# Translational landscape in tomato revealed by transcriptome assembly and ribosome profiling

**DOI:** 10.1101/534677

**Authors:** Hsin-Yen Larry Wu, Gaoyuan Song, Justin W Walley, Polly Yingshan Hsu

## Abstract

mRNA translation is a critical step in gene expression, but our understanding of the landscape and control of translation in diverse crops remains lacking. Here, we combined *de novo* transcriptome assembly and ribosome profiling to study global mRNA translation in tomato roots. Taking advantage of the 3-nucleotide periodicity displayed by translating ribosomes, we identified 354 novel small ORFs (sORFs) translated from previously unannotated transcripts, as well as 1329 upstream ORFs (uORFs) translated within the 5’ UTRs of annotated protein-coding genes. Proteomic analysis confirmed that some of these novel uORFs and sORFs generate stable proteins *in planta*. Compared with the annotated ORFs, the uORFs use more flexible Kozak sequences around translation start sites. Interestingly, uORF-containing genes are enriched for protein phosphorylation/dephosphorylation and signaling transduction pathways, suggesting a regulatory role for uORFs in these processes. We also demonstrated that ribosome profiling is useful to facilitate the annotation of translated ORFs and noncanonical translation initiation sites. In addition to defining the translatome, our results revealed the global control of mRNA translation by uORFs and microRNAs in tomato. In summary, our approach provides a high-throughput method to discover unannotated ORFs, elucidates evolutionarily conserved translational features, and identifies new regulatory mechanisms hidden in a crop genome.

**Significance:** Several studies have shown that altering mRNA translation is a powerful way of improving crop performance. However, due to limited genomic resources and methods, translational regulation remains poorly understood in crops. By leveraging *de novo* transcriptome assembly and ribosome profiling, we mapped and quantified translating ribosomes across the entire transcriptome in tomato roots. This is the first experiment-based survey to systematically identify actively translated ORFs in a crop. Our results reveal numerous unannotated translation events and uncover new regulatory mechanisms of gene expression in tomato. Our approach not only facilitates our understanding of the tomato translational landscape but also provides a practical strategy to study the translatomes of other species.

- The raw RNA-seq and Ribo-seq data have been deposited in the Gene Expression Omnibus (GEO) database under accession no. GSE124962.
- Proteomics raw data files and MaxQuant Search results have been deposited at the MassIVE repository with dataset identifier: MSV000083363.

## Introduction

Besides being an essential step in gene expression, mRNA translation directly shapes the proteome, which contributes to cellular structure, function, and activity in all organisms. The characterization of translational regulation has enabled crop improvement, including increasing tomato sweetness, rice immunity and lettuce resistance to oxidative stress (1–3). However, due to limited genomic resources and methods, most crop translatomes remain understudied.

Ribosome profiling, or Ribo-seq, has emerged as a high-throughput technique to study global translation (4–6). In a Ribo-seq experiment, ribosomes in the sample of interest are immobilized, and the lysate is treated with nucleases to obtain ribosome-protected mRNA fragments (i.e. ribosome footprints). Finally, sequencing of the ribosome footprints reveals the quantity and positions of ribosomes on a given transcript. Features corresponding to active translation, such as 3-nucleotide (nt) periodicity, that emerge from the Ribo-seq data allow novel translation events to be discovered (7–11). For example, upstream open reading frames (uORFs) in the 5’ leader sequence or 5’ untranslated region (UTR) have been shown to be widespread in many protein-coding genes in humans, mouse, zebrafish, yeast, and plants (10– 15). Several well-characterized examples and global analyses indicate that uORFs can modulate the translation of their downstream main ORFs (12, 13, 15–17). Moreover, numerous presumed non-coding RNAs have been found to possess translated small ORFs (sORFs), usually below 100 codons (7, 11, 18, 19). The protein products of these sORFs are so small that they may serve as signaling peptides (16, 19). Despite their importance, uORFs and sORFs are often missing in annotations because computational predictions often assume that 1) protein-coding sequences encode proteins greater than 100 amino acids, and 2) only the longest ORF in a transcript is translated (20, 21). Thus, ribosome profiling provides an unparalleled opportunity to experimentally identify translated ORFs genome-wide in an unbiased manner.

In plant research, ribosome profiling has been used to study the translational regulation in diverse aspects of plant physiology, for example, photomorphogenesis, chloroplast differentiation, cotyledon development, hypoxia, hormone responses, nutrient deprivation, drought, pathogen responses, and biogenesis of small interfering RNAs (15, 18, 22–29). We previously optimized the resolution of this technique, specifically to 3-nt periodicity, to uncover unannotated ORFs in Arabidopsis (11). Although it is clear that ribosome profiling is powerful for studying translational regulation and discovering new translational events in plants, to date, this technique has only been developed in Arabidopsis, soybean, and maize.

Tomato is the most widely cultivated vegetables worldwide (30). It belongs to the Solanaceae, whose members produce important foods, spices, and medicines. Like other crops, tomato has limited genomic resources and optimized methods. For instance, the latest annotation, ITAG3.2 for the ‘Heinz 1706’ cultivar, only contains predicted protein-coding genes while non-coding RNAs and uORFs are not included (31). Here, we performed ribosome profiling, as well as *de novo* transcriptome assembly, to discover non-coding RNAs, uORFs and sORFs, and chart the translational landscape in tomato roots. The mapping and quantification of ribosome footprints in tomato not only uncovered numerous unannotated translation events but also revealed global features involved in translational regulation.

## Results

### Establishment of an experimental and data analysis pipeline to map the tomato translatome

To map actively translated ORFs, we isolated the roots of tomato seedlings (*Solanum lycopersicum*, ‘Heinz 1706’ cultivar) and performed strand-specific RNA-seq and Ribo-seq in parallel (Fig. 1A and 1B). RNA-seq reveals transcript identity and abundance, while Ribo-seq maps and quantifies ribosome occupancy on a given transcript (5). We adapted our protocol and pipeline for Arabidopsis (11) with two major modifications: 1) we increased the amount of RNase I used in tomato ribosome footprinting to achieve comparable resolution (see Materials and Methods for details), and 2) we performed paired-end 100-bp RNA-seq followed by reference-guided *de novo* transcriptome assembly to capture transcripts missing from the ITAG3.2 reference annotation (Fig. 1C, see Materials and Methods for details). This strategy allowed us to map the translated regions in both the annotated and novel transcripts in an unbiased manner using an ORF-finding tool, RiboTaper (8).

**Figure 1:**
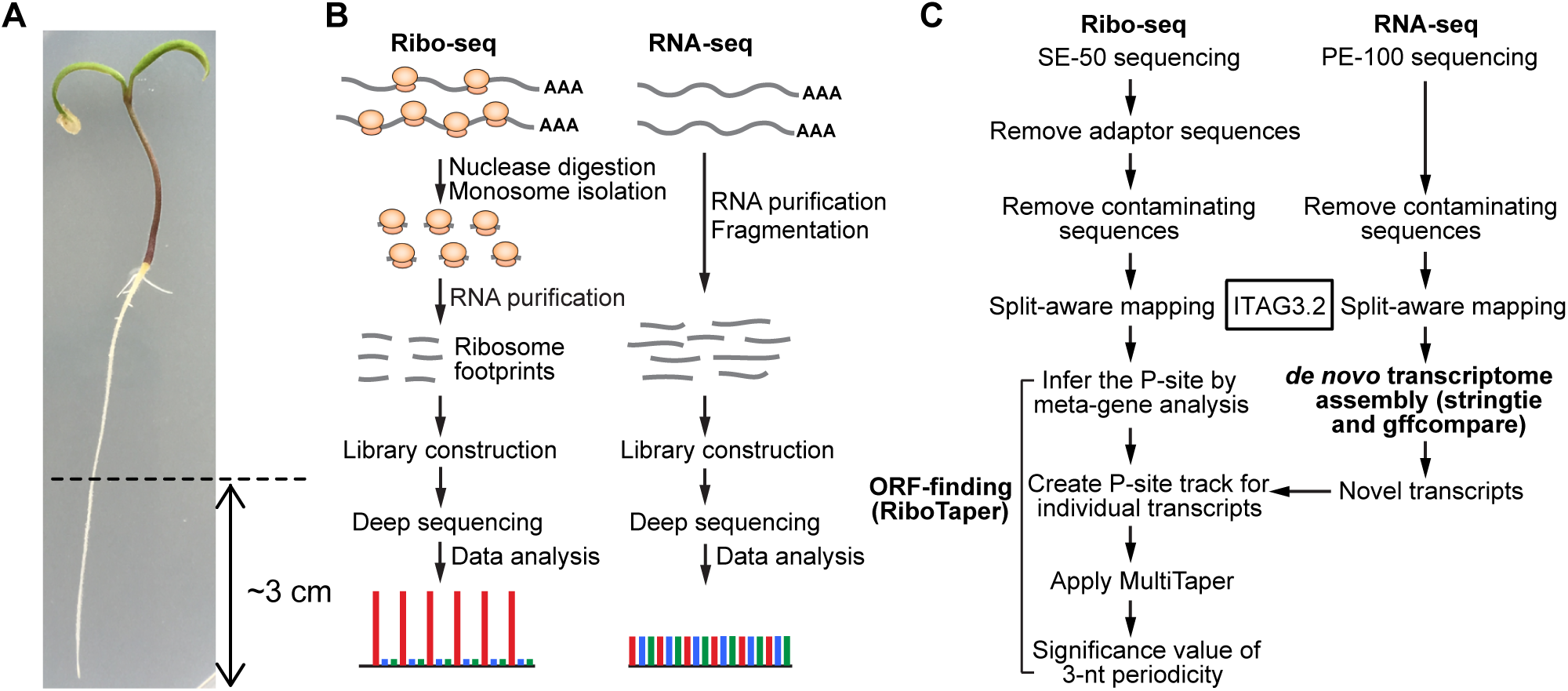
Experimental and data analysis procedures for ribosome profiling in tomato roots. (A) Four-day-old tomato roots (approximately 3 cm from the tip) were used in this study. (B) Experimental workflow for RNA-seq and Ribo-seq and the schematics of their expected read distributions in the three reading frames. (C) Data analysis workflow for reference-guided *de novo* transcriptome assembly and ORF discovery using RiboTaper.

As the quality of ribosome footprints is critical for finding ORFs (11), we first systematically evaluated the Ribo-seq results by mapping the reads to the ITAG3.2 annotation. Consistent with observations in other non-plant organisms and Arabidopsis (4, 7, 11), the dominant ribosome footprints in tomato were 28 nt long (Fig. 2A). Moreover, in contrast to RNA-seq, the Ribo-seq reads predominantly mapped to the annotated coding sequences (CDSs) and were sparse in the 5’ UTRs and 3’ UTRs (Fig. 2B and 2C). The three biological replicates were highly correlated, as indicated by the Pearson correlation, in both Ribo-seq (r = 0.998~1) and RNA-seq (r = 0.998~0.999) (Fig. S1A and B). Overall, the RNA-seq and Ribo-seq datasets also showed a strong positive correlation (Pearson correlation after removing two extreme outliers, r = 0.878-0.880; Spearman correlation ρ = 0.912-0.915) (Fig. S1C-F). Most importantly, the distribution of ribosome footprints within the CDS displayed clear 3-nt periodicity, a signature of translating ribosomes, which decipher 3 nt at a time (Fig. 2C and Fig. S2). Analyzing the distribution of footprints relative to the annotated translation start/stop sites allowed us to infer that the codon at the P-site within the ribosome is located between the 13^th^ and 15^th^ nts for 28-nt footprints, and so on for specific footprint lengths (Fig. S2-S3). To visualize the position of the codon being translated, hereafter, we use the first nucleotide of the P-sites (denoted as P-site signals) to indicate the positions of the footprints on the transcripts (Fig. 2C). The robustness of the 3-nt periodicity can be quantified based on the percentage of reads in the expected reading frame (shown in red in Fig. 2C and hereafter). At a global level, our 28-nt footprints resulted in 85.5% in-frame reads. Together, these results demonstrate that our tomato Ribo-seq dataset is of high quality compared to datasets from plants and other organisms (7, 11, 32–34).

**Figure 2:**
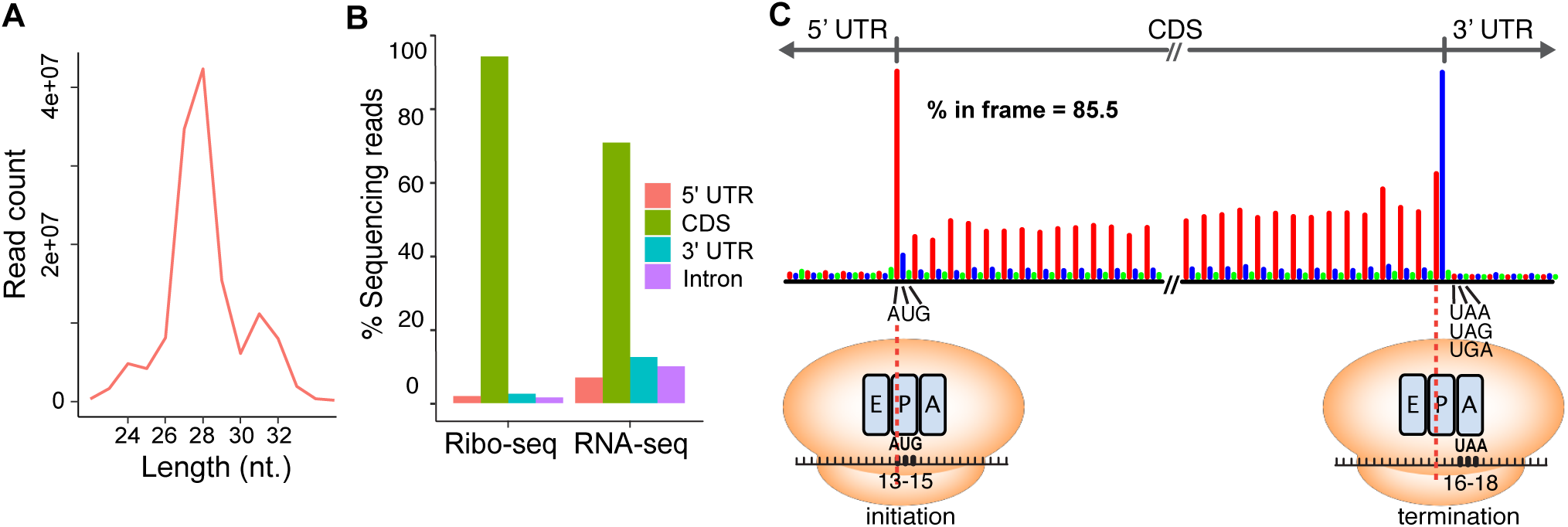
Ribosome footprints are enriched in coding sequences and display strong 3-nt periodicity. (A) The distribution of read length of the ribosome footprints. (B) The distribution of the Ribo-seq and RNA-seq reads in different genomic features annotated in ITAG3.2. (C) Meta-gene analysis of the 28-nt ribosome footprints near the annotated translation start and stop sites defined by ITAG3.2. The red, blue and green bars represent reads mapped to the first (expected), second and third reading frames, respectively. The majority of footprints were mapped to the CDS in the expected reading frame (85.5% in frame). For each read, only the first nucleotide in the P-site was plotted (for details, see Figs. S2-S3). The A-site (<u>a</u>minoacyl-tRNA entry site), P-site (<u>p</u>eptidyl-tRNA formation site) and E-site (uncharged tRNA <u>e</u>xit site) within the ribosomes at translation initiation and termination, and the inferred P-site (13^th^-15^th^ nts) and A-site (16^th^-18^th^ nts) are illustrated. The original meta-plots generated by RiboTaper for all footprint lengths are shown in Fig. S2.

Next, we performed reference-guided *de novo* transcriptome assembly for the RNA-seq data using *stringtie*, a transcript assembler (35). Then, the newly assembled transcriptomes from the replicates were merged and compared to the ITAG3.2 annotations using *gffcompare* (36) software (Fig. 1C). In total, we uncovered 2263 unannotated transcripts that could potentially encode for novel proteins. These transcripts could be classified into six groups based on their strands and genomic positions relative to existing gene features, such as intergenic (class-u), cis-natural antisense transcripts (cis-NAT, class-x), intronic (class-i) and others (class-y and class-o) (Fig. 3A and 3C, the nomenclature and descriptions of novel transcripts are adapted based on the *gffcompare* software (36)). Class-s is expected to result from mapping errors (36) and was included in our downstream analysis as a negative control. The most abundant novel transcripts in our data were intergenic transcripts (class-u; 1260) and cis-NATs (class-x; 568). All six classes of novel transcripts, along with the annotated genes in ITAG3.2, were used to find translated ORFs.

**Figure 3:**
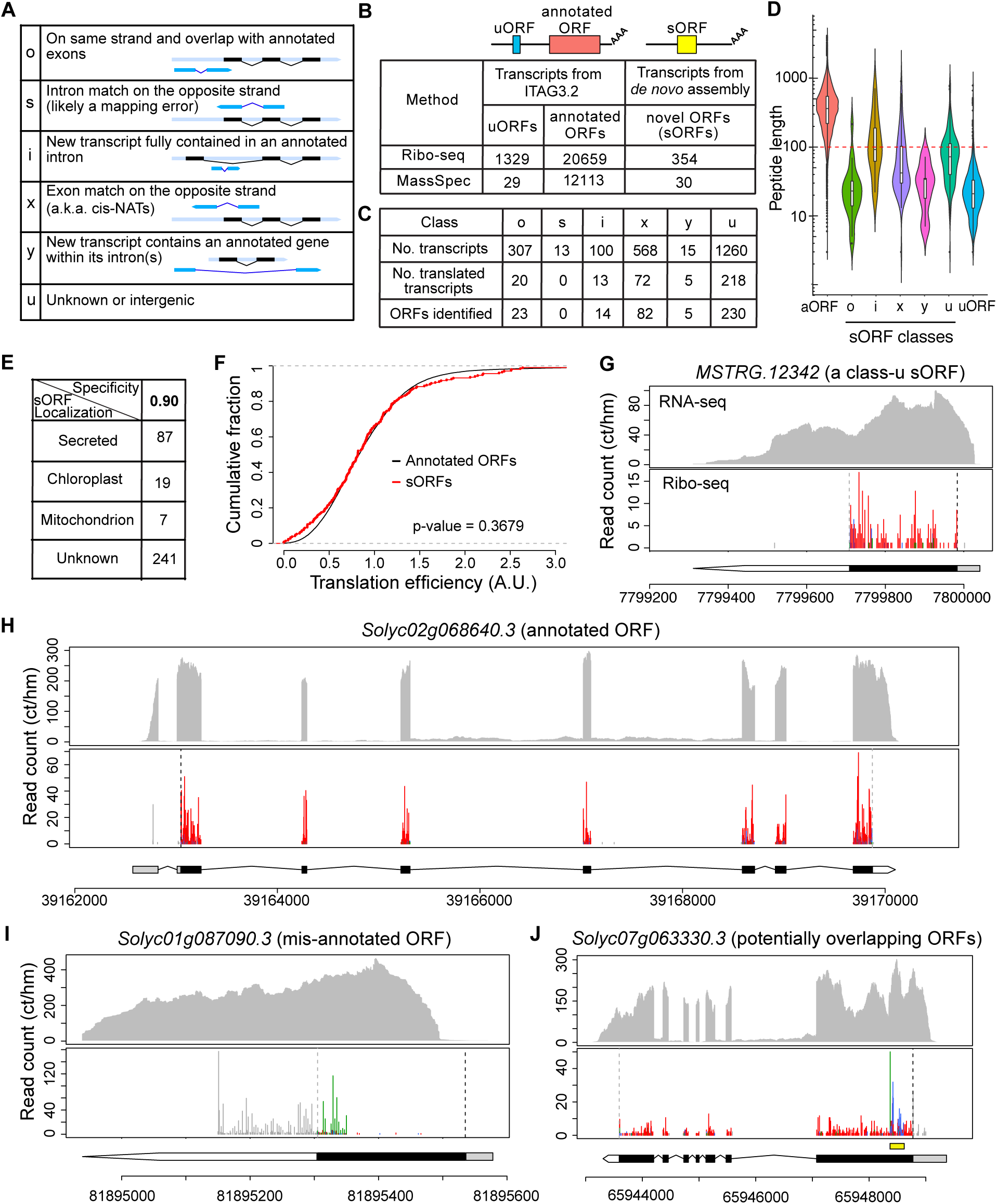
The translational landscape of the tomato root. (A) Classes of newly assembled transcripts identified by stringtie and gffcompare and used in downstream ORF identification. This figure was adapted from the gffcompare website (36). (B) Summary of translated ORFs identified by RiboTaper in our dataset and peptide support from mass spectrometry data. The uORFs and annotated ORFs were identified from the 5' UTRs and expected CDSs of annotated protein-coding genes in ITAG3.2, respectively. The novel ORFs were identified from the newly assembled transcripts. The bottom row indicates the number of proteins in each category supported by mass spectrometry datasets, either from our own proteomic analysis or searches against publicly available data. (C) Summary of newly assembled transcripts and ORFs identified in each class of newly assembled transcripts. The total number of transcripts, number of transcripts identified as translated and the total number of translated ORFs are listed. (D) Size distribution of each class of sORFs, uORFs and annotated ORFs (aORFs). (E) Predicted subcellular localization of proteins encoded by the sORFs. The prediction was performed using TargetP (85) with specificity 0.9 as a cutoff. (F) Translation efficiency of sORFs compared with annotated ORFs. Only the coding regions were used to compute the TPM and translation efficiency of each transcript. For the x-axis, only the range from 0 to 3 (arbitrary unit) is shown. A two-sample Kolmogorov-Smirnov test was used to determine statistical significance. (G-J) RNA-seq coverage and Ribo-seq periodicity in different genes: (G) an intergenic sORF on chromosome 4; (H) an annotated coding gene that has good support from the Ribo-seq data for the predicted gene model; (I) a mis-annotated ORF; note the Ribo-seq reads do not match the CDS in the gene model and a different reading frame is used; (J) a transcript with a potentially overlapping ORF within the annotated ORF. In (G-J), the x-axis indicates the genomic coordinate of the gene. The y-axis shows the normalized read count (counts per hundred million reads). Ribo-seq reads are shown by plotting the first nucleotide of their P-sites (denoted as the P-site signals). The black and gray dashed vertical lines mark the predicted translation start and stop sites, respectively. The red, blue and green lines in the Ribo-seq plot indicate the P-site signals mapped to the first (expected) reading frame and the second and third reading frames, respectively. Hence, a higher ratio of red means better 3-nt periodicity. For the gene model beneath the Ribo-seq data, the gray, black and white areas indicate the 5' UTR, CDS and 3' UTR, respectively. In (J), the yellow box above the gene model indicates the region with a potential ORF overlapping with the annotated ORF.

### Translational landscape of tomato roots as defined by ribosome profiling

After collecting the transcript information, we used RiboTaper (8) to interrogate both the annotated transcripts in ITAG3.2 and the newly assembled transcripts to search for all possible ORFs in the transcriptome. RiboTaper examines the P-site signals at each possible ORF and tests whether the signals display a statistically significant 3-nt periodicity (8). As a quality control, we first examined number of translated ORFs detected at annotated coding regions. In total, 20659 annotated ORFs were identified as translated in our dataset (Fig. 3B and Dataset_S1A). Among 20285 annotated protein-coding transcripts that have reasonable expression levels (transcript per million (TPM) > 0.5 in RNA-seq), 18626 (92%) have translated ORFs identified. This indicates our approach to identifying translated ORFs is efficient and robust. In addition to annotated ORFs, there were 1329 uORFs translated upstream of the annotated ORFs within the 5’ UTR (Fig. 3B, Dataset_S1B and Dataset_S3). Notably, since only approximately half of the transcripts in ITAG3.2 (17684 out of 35768) have an annotated 5’ UTR, and because RiboTaper can only identify ORFs in defined transcript ranges, the total number of uORFs in tomato root is clearly an underestimate. Excitingly, we identified 354 novel translated ORFs from the newly assembled transcripts (Dataset_S1C and Dataset_S4). These novel ORFs were found in different classes of transcripts, but none were detected in the negative control, class-s (Fig. 3C). As expected, most of the novel ORFs were relatively small; ~71% of them encode proteins of less than 100 a.a. (Fig. 3D). Due to their relatively small size, hereafter, we call them <u>s</u>mall <u>ORFs</u> (sORFs). The average lengths of the uORFs, sORFs, and annotated ORFs are 31, 95 and 422 amino acids, respectively. Among the 354 sORFs, 87 have a predicted signal peptide and are expected to be secreted proteins/peptides (Fig. 3E and Dataset_S1D). To test if the sORFs and annotated ORFs have similar translational properties, we compared their translation efficiency (see the definition in the Materials and Methods) and found that they were statistically indistinguishable (Fig. 3F). This result supports the newly identified sORFs are genuine protein-coding genes in the tomato genome.

The majority of the identified ORFs have high fractions of P-site signals mapped to the expected reading frame (Fig. S4). Visualizing the profiles of individual transcripts confirmed that both the sORFs and numerous annotated ORFs display strong 3-nt periodicity within the identified coding regions (Fig. 3G-H). Therefore, by combining the high-quality Ribo-seq data with RiboTaper analysis, we not only validated many of the annotated gene models but also discovered new ORFs.

### sORFs are evolutionarily conserved

Previously, we identified 27 sORFs in Arabidopsis by applying RiboTaper on Ribo-seq data (11). Eight of the Arabidopsis sORFs have known tomato homologs. Our tomato root data showed that seven of the conserved sORFs were both transcribed and translated (Fig. S5A-D). Since Arabidopsis and tomato diverged approximately 100 million years ago (37), our data support that some sORFs are conserved across evolution.

If the sORFs encode functionally important proteins, we would expect them to be evolutionarily conserved across a wide range of genomes. We performed tblastn using 157 sORFs that were 16~100 a.a. long on 10 diverse plant genomes, including a wild tomato (*S. pennellii*), potato (belongs to the same family as tomato, the Solanaceae), four dicots in other families, two monocots, a lycophyte and a moss (Fig. S6). In total, we found 139 sORFs have homologs in at least one other plant genome. Almost all have homologs within the Solanaceae, often in both wild tomato and potato and with high amino acid identity. Some of the sORFs are highly conserved in all 10 genomes (Fig. S6). Importantly, the conserved patterns among the homologs correlate well with their phylogenic relationships, indicating that these sORF homologs are unlikely to be false positives that randomly occurred in the blast search. Taken together, these results suggest the functional significance of these sORFs throughout evolution.

### Some sORFs and uORFs generate stable proteins *in planta*

To evaluate whether the novel ORFs, including sORFs and uORFs, accumulate stable proteins *in planta* and to validate our Ribo-seq results, we performed a proteogenomic analysis (38) to identify “novel” peptides arising from these unannotated ORFs. Because the sORFs and uORFs are quite small, their protein products do not always generate peptides with ideal size and/or mass-to-charge ratios that are suitable for detection by mass spectrometry (MS). To increase the diversity of peptides for MS analysis, we extracted proteins from the roots and shoots of tomato seedlings and digested the proteins into peptides using trypsin or GluC, independently, prior to two-dimensional liquid chromatography-tandem mass spectrometry (2D-LC-MS/MS). As the sORFs and uORFs are currently missing from the tomato annotation, we created a custom protein database derived from our Ribo-seq data to assist in identifying these unannotated proteins. In addition, we used our custom protein database to search publicly available proteomic data from the tomato fruit (ProteomeXchange PXD004887) and pericarp (ProteomeXchange PXD004947) (39, 40). In total, we identified 12172 proteins, including 29 sORFs and 30 uORFs, with at least one unique peptide from these six proteomic datasets (Fig. 3B, Dataset_S1E-F and Dataset_S2A-C). Despite the limitations of MS in small protein identification, our results support that some uORFs and sORFs accumulate stable proteins *in planta*.

### Ribo-seq fine-tunes and improves genome annotation

Comparing the RiboTaper output and the annotated gene models, we found cases in which the translated ORFs were dramatically different from the predicted gene models. For example, translation may occur in a different reading frame or at a distinct region on the transcript (Figs. 3I and S7). Thus, Ribo-seq provides a high-throughput experimental approach to validate and improve genome annotation. Furthermore, in several cases, using visual inspection, we found regions that appear to contain a short ORF that overlaps with the long annotated ORF but uses a different reading frame (e.g., Fig. 3J). These overlapping ORFs are reminiscences of non-upstream coding ORFs identified in human genome (41), and their functional importance is still unknown.

The translation start sites in the genome annotation are typically defined computationally, and often the most upstream AUG is predicted to be the start codon. Unexpectedly, in 64 genes, the RiboTaper-defined translation start sites were actually upstream of the annotated start sites (e.g., Fig. 4A and Dataset_S1G). In contrast, some ORFs appeared to use start sites downstream of the annotated start sites (e.g., Figs. 4B and S5B). Currently, ITAG3.2 contains only one isoform per gene, and hence only one transcription start site is predicted per gene. It is possible that in some cases, translation starts downstream of the annotated site because transcription initiates downstream of the annotated transcription start site. Nonetheless, it appears that the most upstream AUG is not always used as the translation start site.

**Figure 4:**
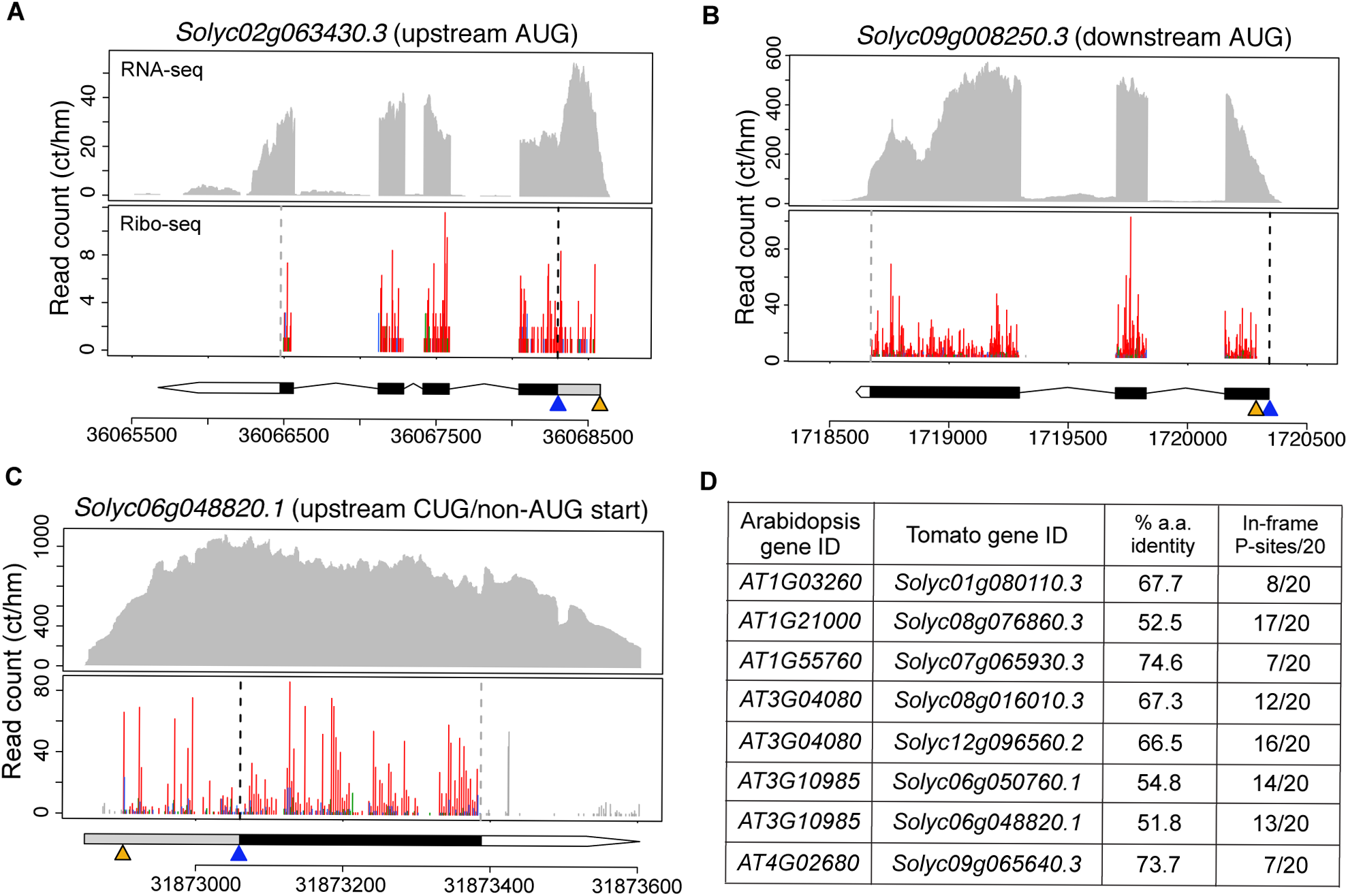
Upstream/downstream start sites and non-AUG start sites. (A-B) Examples of the usage of an upstream start site (A) or a downstream start site (B). The gene model and data presentation are the same as those described in the legend of Fig. 3. The blue triangle marks the location of the annotated translation start site. The orange triangle marks the location of the RiboTaper-identified translation start site. (C) A tomato homolog of an Arabidopsis gene that was predicted to use an upstream CUG start site (orange triangle). Note the abundant in-frame P-site signals upstream of the annotated AUG start (blue triangle) in the 5’ UTR. (D) Conservation of potential CUG/non-AUG start sites. The Arabidopsis gene ID, tomato gene ID, percent amino acid identity, and number of in-frame P-site positions with Ribo-seq reads within the first 20 codons upstream of the AUG in our tomato root data are shown.

Non-AUG translation initiation has been discovered in animals and plants (42–45). Twelve evolutionarily conserved noncanonical translation starts upstream of the most likely AUG have been predicted in Arabidopsis (43), and we previously showed that at least one of them, in *AT3G10985*, has high Ribo-seq coverage using a CUG codon (11). The profile of the tomato homolog of *AT3G10985* confirmed the possible usage of the CUG start site (Fig. 4C). Next, we identified tomato homologs of all twelve predicted noncanonical-start genes and systematically checked their Ribo-seq coverage upstream of the annotated AUG start sites. We selected genes that met the following criteria: 1) the Ribo-seq reads have at least 7 in-frame P-site positions within the first 20 codons upstream of the AUG, and 2) there is no stop codon within the first 20 codons upstream of the AUG. We found eight tomato genes that met the above criteria contain abundant reads upstream of the annotated AUG, suggesting that they use non-AUG start sites (Fig. 4D). Although Arabidopsis and tomato diverged approximately 100 million years ago (37), the usage of noncanonical translation initiation remains conserved in these homologs.

### uORFs regulate translation efficiency

Using RiboTaper, we identified 1329 translated uORFs based on their significant 3-nt periodicity (Fig. 3B, Dataset_S1B and Dataset_S3). These uORFs included previously predicted conserved uORFs in the tomato *SAC51* homolog (Fig. 5A)(46), as well as novel uORFs in numerous coding genes (e.g. Fig. 5B). Manual inspection of these transcripts suggested that the high stringency of RiboTaper might miss uORFs with lower periodicity, overlapping uORFs and non-AUG-start uORFs. For example, the second of the three uORFs in the tomato *SAC51* transcript (Fig. 5A) was not identified as coding by RiboTaper, presumably due to the imperfect periodicity in this area. Nevertheless, those identified are high-confidence translated uORFs.

**Figure 5:**
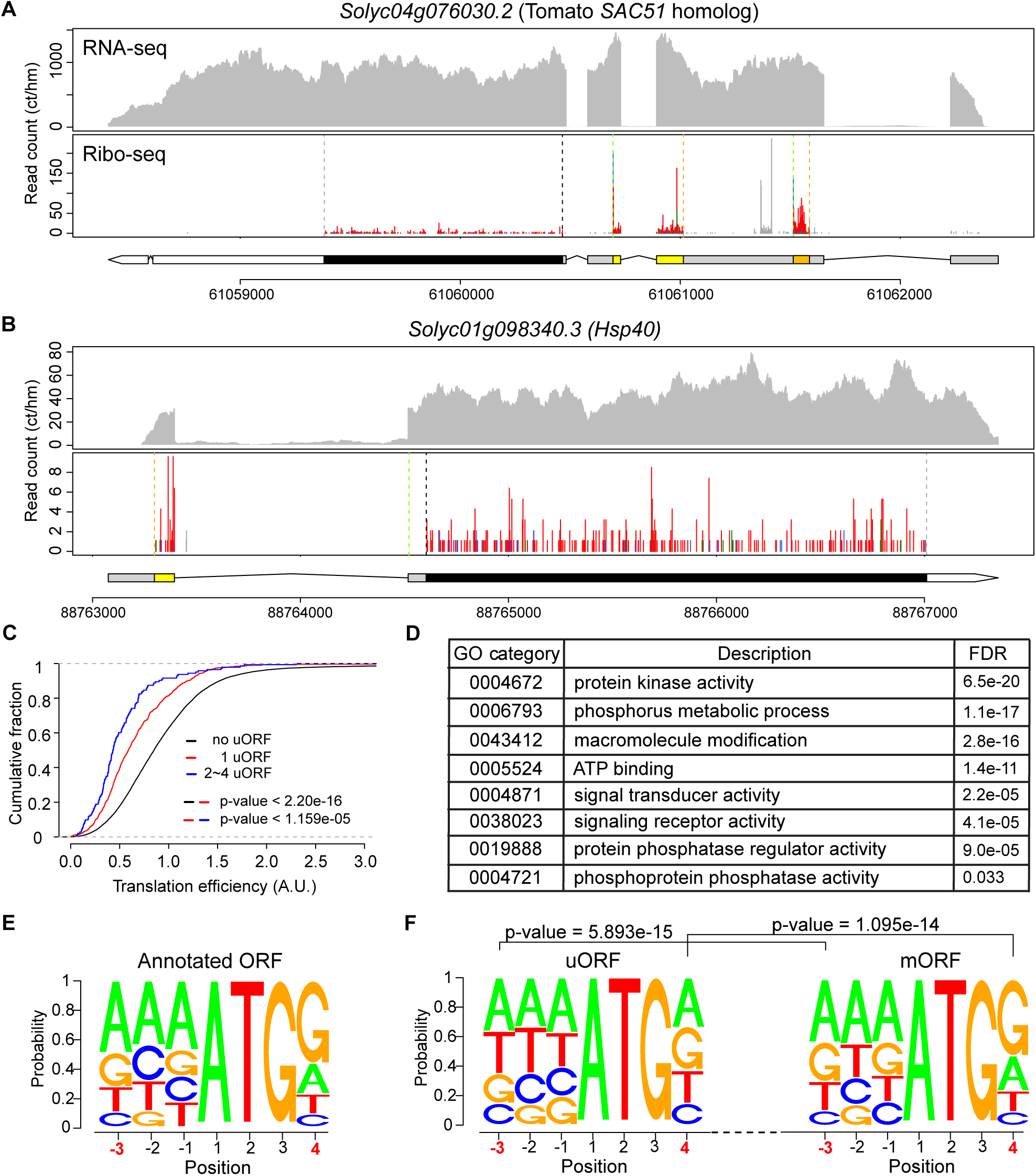
uORFs repress translation efficiency of their downstream main ORFs and contain less-pronounced Kozak sequences. (A, B) Profiles of genes containing conserved uORFs (A) or a novel uORF (B). The gene model and data presentation are the same as those described in the legend of Fig. 3. The uORFs are labeled with yellow and orange boxes in the gene models. For the uORFs, the orange and green dashed vertical lines mark the translation start and stop sites, respectively. (C) The translation efficiency (TE) of the main ORFs for transcripts containing a different number of translated uORFs. Only the coding regions were used to compute the TPM and translation efficiency of each transcript. The colored bars before the p-values indicate the pairs of data used to determine statistical significance. The p-values were determined with two-sample Kolmogorov-Smirnov tests. (D) Selected non-redundant GO categories for genes containing one or more uORFs. (E, F) Kozak sequences of annotated ORFs, uORFs, and uORF-associated main ORFs. The statistical significance in (F) was determined using Chi-squared tests.

Several studies have reported that uORFs repress the translation of their downstream main ORFs (12, 13, 15–17). Consistent with these reports, we found that globally, transcripts containing uORFs have lower translation efficiency than those without uORFs (Fig. 5C). In addition, more uORFs in a transcript correlate with stronger translational repression (Fig. 5C). To investigate which physiological pathways might be regulated by uORFs, we checked the gene ontology (GO) terms of the uORF-containing genes. Intriguingly, uORF-containing genes were enriched for protein kinases and phosphatases, as well as signal transduction (Fig. 5D), implying that a substantial portion of the protein phosphorylation/dephosphorylation and signal transduction pathways in tomato are likely translationally regulated through uORFs.

Translation start sites have a well-defined Kozak consensus sequence in different organisms (47, 48). For example, the conserved nucleotides at positions −3 and +4 of the Kozak sequence in plants are purines (A/G) and G, respectively (48). As expected, we observed this conserved pattern among the annotated ORFs (Fig. 5E). Next, we examined the Kozak consensus sequences of the translated uORFs and their downstream main ORFs. While the downstream main ORFs also favor the conserved nucleotides at −3 and +4 of the Kozak sequence, this pattern is missing in the uORFs (Fig. 5E and 5F). Similar results were observed in the Kozak sequences of the uORFs and downstream main ORFs in Arabidopsis (15). The poorly conserved Kozak sequences might allow for more-flexible translation initiation and regulation of uORFs.

### Regulation of gene expression by microRNAs

MicroRNAs regulate gene expression through mRNA cleavage and translational repression (49, 50). The roles of microRNAs in tomato are less well understood than in Arabidopsis. We first predicted 6312 microRNA target genes in tomato (Dataset_S1H-I) using psRNATarget (51). Next, we compared their RNA-seq and Ribo-seq levels and the translation efficiency of the microRNA targets and other coding genes globally. The transcript levels of the miRNA targets were slightly but significantly reduced, consistent with the possibility that microRNAs regulate gene expression through mRNA cleavage (Fig. 6A). In addition, both the Ribo-seq levels and translational efficiency of the microRNA target genes were reduced (Fig. 6B and 6C), consistent with prior observations of translational repression mediated by microRNAs (52). Thus, our results suggest that globally, microRNAs regulate gene expression at both the transcriptional and translational levels in tomato.

**Figure 6:**
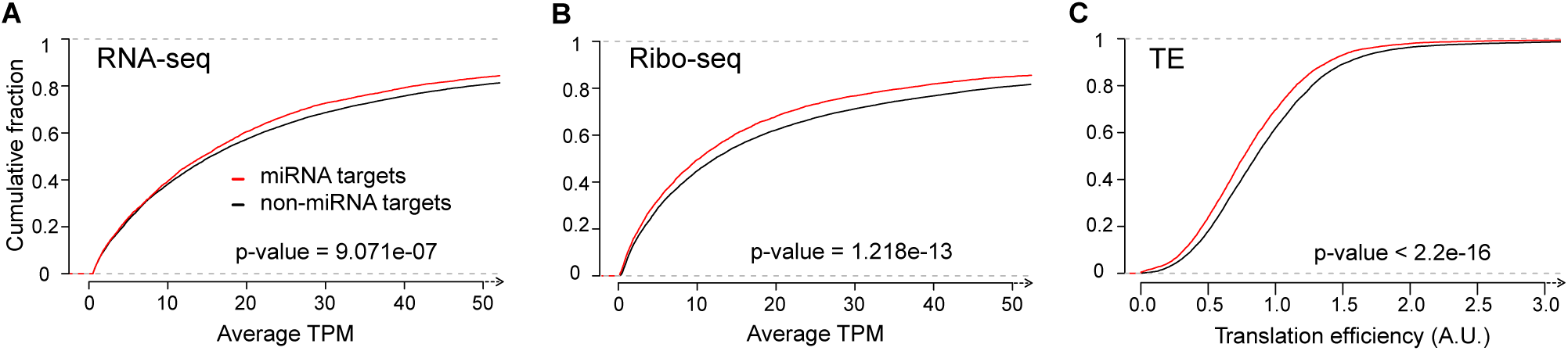
Regulation of gene expression by microRNAs (miRNAs). Cumulative distributions of (A) RNA-seq, (B) Ribo-seq and (C) translation efficiency (TE) of miRNA targets and non-miRNA target genes. For the x-axis in (A) and (B), only the range from 0 and 50 (TPM) is shown. Only the coding regions were used to compute the TPM and translation efficiency of each transcript. The p-values were determined with two-sample Kolmogorov-Smirnov tests.

## Discussion

In our previous work in Arabidopsis, we systematically tested the lysis buffers used in ribosome profiling and identified one that would yield optimal 3-nt periodicity (11). As different plant materials require different amounts of the RNase I for digestion (11), we found increasing the RNase I but using the same buffer we developed in Arabidopsis yielded comparable resolution in tomato. Overall, we observed similar features between our Arabidopsis and tomato Ribo-seq data, including a) the most abundant ribosome footprints being 28 nt long (Fig. 2A), b) the inferred P-site being located between the 13^th^ and 15^th^ nt in 28-nt footprints (Figs. 2C and S2-S3), c) globally, footprints that are 28 nt or fewer displaying better 3-nt periodicity (Fig. S2), and d) RNase I preferentially digesting the 5’ end of the ribosome footprints when the footprints are shorter than 28 nt (Figs. S2-S3).

We identified a higher number of novel translation events, including uORFs and sORFs, in tomato than in Arabidopsis. Two factors may have contributed to this difference. The first is the sample harvesting method. Our Arabidopsis seedlings were grown hydroponically; thus, we had to blot the plants with paper towels to remove the liquid media before freezing the tissue with liquid nitrogen (11). In contrast, the tomato seedlings were grown vertically on solid media in Petri dishes, and the roots grew along the surface of the media. Therefore, the harvest time was significantly shorter for tomato than for Arabidopsis. Translation is an extremely dynamic process, as 5~20 codons can be translated per second (53, 54). It is recommended to have prompt sample harvesting and freezing in liquid nitrogen to preserve the translational status (55). If some ribosomes run off during sample harvest, the shorter ORFs, such as uORFs and sORFs, are the most likely to be affected by the harvesting time. The second possible explanation for the different numbers of novel ORFs found in Arabidopsis and tomato is the genome annotations. The Arabidopsis genome has been extensively studied. Many small peptides have been computationally predicted and experimentally characterized (16, 56–61); thus, unannotated sORFs are less likely to be found in Arabidopsis. In our previous work, we searched sORFs among the annotated non-coding RNAs in TAIR10 (11, 62). Unlike Arabidopsis, the latest tomato annotation does not contain predicted non-coding RNAs (31).

Instead of using available transcript information, we performed transcriptome assembly to identify transcripts missing in the annotation to uncover novel ORFs. As a result, we could search ORFs in all transcripts expressed in our tomato samples, but in Arabidopsis, we only searched for known transcripts. Both the less-complete peptide annotation and the transcriptome assembly may have contributed to the higher number of sORFs discovered in tomato.

Ribo-seq has been integrated into proteomic research to achieve deeper proteome coverage (8, 63–65). Unlike DNA or RNA molecules, which can be sequenced using genomic technologies, proteins are typically identified by matching MS spectra to theoretical spectra from candidate peptides in a reference protein database. Before ribosome profiling became available, to include potential protein sequences, the conventional proteogenomics approach used either three-frame-translation using transcriptome data or six-frame-translation using genomic sequences (38, 66). Integrating Ribo-seq data into the construction of protein databases for proteogenomic studies has several advantages: 1) Ribo-seq discovers unannotated translation events and thus enables the identification of novel proteins that were previously missed in the annotation, and 2) compared with three-frame or six-frame translation, Ribo-seq reduces the search space and false positives. Therefore, our custom protein database, built based on the Ribo-seq data, may aid in proteomic research in tomato.

Several studies have illustrated the power of altering mRNA translation via uORFs to improve agriculture (1–3). For example, engineering rice that specifically induces defense proteins when a uORF is repressed by pathogen attack enables immediate plant resistance without compromising plant growth in the absence of pathogens (2). The identification of translated ORFs provides new possibilities to fine-tune the synthesis of proteins involved in diverse physiological pathways. Notably, the number of uORFs in tomato is still an underestimate. Approximately half of tomato genes still lack annotated 5’ UTRs, and RiboTaper only searches for potential translated ORFs in defined transcript regions. Thus, uORFs could be an even more widespread mechanism to control translation in tomato.

Peptide signaling is crucial for cell-cell communication in numerous aspects of plant development and stress responses (16, 61). We found 87 sORFs that encode potential secreted peptides. However, as about 50% of secreted proteins in plants lack a well-defined signal peptide (67), some sORFs without a predicted signal peptide may still be secreted. In addition, sORF products without a signal peptide have been found to play an important role in a wide range of physiological processes in plants, such as vegetative and reproductive development, siRNA biogenesis, and stress tolerance (68–72). Therefore, the identification of sORFs using ribosome profiling facilitates potential applications of these peptides in improving crop performance.

In conclusion, our approach combining transcriptome assembly and ribosome profiling enabled identification of translated ORFs genome-wide in tomato and revealed conservation of translational features across evolution. Our results not only provide valuable information to the plant community but also present a practical strategy to study translatomes in other less-well annotated organisms.

## Materials and Methods

### Plant materials and preparation of lysates for RNA-seq and Ribo-seq

Tomato seeds (*Solanum lycopersicum*, ‘Heinz 1706’ cultivar) were obtained from the C.M. Rick Tomato Genetics Resource Center (Accession: LA4345) and bulked. For each replicate, ~300 tomato seeds were surface-sterilized in 70% ethanol 5 minutes followed by bleach solution (2.4% NaHClO, 0.3% Tween-20) for 30 minutes with shaking. The seeds were then washed with sterile water 5 times. Next, the seeds were stratified on 1x Murashige and Skoog media (4.3 g/L Murashige and Skoog salt, 1% sucrose, 0.5 g/L MES, pH 5.7, 1% agar), and kept at 22 °C in the dark for 3 days before being grown under 16-hour light/8-hour dark conditions at 22°C for 4 days. Seedlings that germinated at approximately the same time and of similar size were selected for the experiments. Roots (~3 cm from the tip) from ~180 plants were harvested at ZT 3 (3 hours after lights on) in batches and immediately frozen in liquid nitrogen. The frozen tissues were pooled and pulverized in liquid nitrogen using a mortar and pestle. Approximately 0.4 g of tissue powder was resuspended in 1.2 mL lysis buffer (100 mM Tris⋅HCl (pH 8), 40 mM KCl, 20 mM MgCl_2_, 2% (vol/vol) polyoxyethylene (10) tridecyl ether (Sigma P2393), 1% sodium deoxycholate (Sigma D6750), 1 mM DTT, 100 μg/mL cycloheximide, and 10 unit/mL DNase I (Epicenter D9905K)) as described in Hsu *et al.* 2016 (11). After incubation on ice with gentle shaking for 10 minutes, the lysate was spun at 4°C at 20,000 x g for 10 minutes. The supernatant was transferred to a new tube and divided into 100-µL aliquots. The aliquoted lysates were flash frozen in liquid nitrogen and stored at −80°C until processing.

### RNA purification and RNA-seq library construction

For RNA-seq samples, 10 µL 10% SDS was added to the 100-µL lysate aliquots described above. RNA greater than 200 nt was extracted using a Zymo RNA Clean & Concentrator kit (Zymo Research R1017). The obtained RNA was checked with a Bioanalyzer (Agilent) RNA pico chip to access the RNA integrity, and a RIN value ranging from 9.2 to 9.4 was obtained for each replicate. Ribosomal RNAs (rRNAs) were depleted using a RiboZero Plant Leaf kit (Illumina MRZPL1224). Next, 100 ng of the rRNA-depleted RNA was used as the starting material, fragmented to ~200 nt based on the RIN reported by the Bioanalyzer, and processed using an NEBNext Ultra Directional RNA Library Prep Kit (NEB E7420S) to create strand-specific libraries. The libraries were barcoded and enriched using 11 cycles of PCR amplification. The libraries were brought to equal molarity, pooled and sequenced on one lane of a Hi-Seq 4000 using PE-100 sequencing.

### Ribosome footprinting and Ribo-seq library construction

The Ribo-seq samples were prepared based on Hsu *et al.* 2016 (11) with modifications described as follows, which optimize the method for tomato. Briefly, the RNA concentration of each lysate was first determined using a Qubit RNA HS assay (Invitrogen Q32852) using a 10-fold dilution. Next, 100 µL of the lysate described above was treated with 100 units of nuclease (provided in the TruSeq Mammalian Ribo Profile Kit, Illumina RPHMR12126) per 40 µg of RNA with gentle shaking at room temperature for one hour. The nuclease reaction was stopped by immediately transferring to ice and adding 15 µL of SUPERase-IN (Invitrogen AM2696). The ribosomes were isolated using illustra MicroSpin S-400 HR columns (GE Healthcare 27514001). RNA greater than 17 nt was purified first (Zymo Research R1017), and then RNA smaller than 200 nt was enriched (Zymo Research R1015). Next, the rRNAs were depleted using a RiboZero Plant Leaf kit (Illumina MRZPL1224). The rRNA-depleted RNA was then separated via 15% (wt/vol) TBE-urea PAGE (Invitrogen EC68852BOX), and gel slices ranging from 28 to 30 nt were excised. Ribosome footprints were recovered from the excised gel slices using the overnight elution method, and the sequencing libraries were constructed according to the TruSeq Mammalian Ribo Profile Kit manual. The final libraries were amplified via 9 cycles of PCR. The libraries were brought to equal molarity, pooled and sequenced on two lanes of a Hi-Seq 4000 using SE-50 sequencing.

### RNA-seq and Ribo-seq data analysis

The raw RNA-seq and Ribo-seq data and detailed mapping parameters have been deposited in the Gene Expression Omnibus (GEO) database (www.ncbi.nlm.nih.gov/geo) under accession no. GSE124962. The tomato reference genome sequence and annotation files used in this study were downloaded from the Sol Genomics Network (31). The adaptor sequence AGATCGGAAGAGCACACGTCT was first removed from the Ribo-seq data using FASTX_clipper v0.0.14 (http://hannonlab.cshl.edu/fastx-toolkit). For both RNA-seq and Ribo-seq, the rRNA, tRNA, snRNA, snoRNA and repeat sequences were removed using Bowtie2 v2.3.4.1 (73). Since ITAG3.2 does not contain rRNA, tRNA, snRNA, or snoRNA information, these sequences were extracted from the SL2.5 genome assembly with the ITAG2.4 annotation (31) for contaminating sequence removal with Bowtie2. The pre-processed RNA-seq and Ribo-seq files were then used to calculate the read distribution in different gene features (Fig. 2B) using the featureCounts function of the Subread package v1.5.3 (74).

Next, the pre-processed RNA-seq and Ribo-seq reads were mapped to the tomato reference genome sequence SL3.0 with the ITAG3.2 annotation using the STAR v2.6.0.c (75). The reference-guided *de novo* assembly of the mapped RNA-seq reads was performed with stringtie v1.3.3b (35), and the newly assembled gtf files were compared to ITAG3.2 using gffcompare v0.10.1 (36). The i, x, y, o, u, s classes of new transcripts (see Fig. 3A for details) and their descriptions were extracted from the gffcompare output gtf and concatenated with ITAG3.2. This combined gtf (referred as “Tomato_Root_ixyous+ITAG3.2.gtf”; submitted to GEO as a processed file within GSE124962) was used to map the RNA-seq and Ribo-seq reads again with STAR. Notably, all six classes of novel transcripts in Tomato_Root_ixyous+ITAG3.2.gtf were assigned as ncRNAs, and this gtf was used for downstream RiboTaper analysis. The three biological replicates of the mapped bam files for RNA-seq were merged into one large bam file with SAMtools v1.8 (76). The three mapped Ribo-seq bam files were also merged. The two merged bam files above were then used for ORF discovery with RiboTaper v1.3 (8).

For RiboTaper analysis, the RiboTaper annotation files and the offset parameters (i.e., the inferred P-site position for each footprint length) were first obtained. The RiboTaper annotation files were generated using the create_annotations_files.bash function in the RiboTaper package using SL3.0 assembly and the Tomato_Root_ixyous+ITAG3.2.gtf. To obtain the offset parameters, the create_metaplots.bash and metag.R functions in the RiboTaper package were used to generate meta-gene plots. The offset parameters were identified through the meta-gene plots. For 24-, 25-, 26-, 27-, 28-nt footprints, the offset values were 8, 9, 10, 11, and 12, respectively (Fig. S3). Next, we performed RiboTaper analysis using the RiboTaper annotation, offset parameters, and RNA-seq and Ribo-seq bam files. The coding sequences identified by RiboTaper from the newly assembled transcripts were extracted from the translated_ORFs_filtered_sorted.bed file and integrated with Tomato_Root_ixyous+ITAG3.2.gtf to generate Dataset_S3_uORF.gtf and Data_sORF.gtf.

We then mapped the Ribo-seq and RNA-seq data again to the CDS ranges with STAR, and the transcripts per million (TPM) for the CDS of each transcript was quantified via RSEM v1.3.0 (77). The formula to calculate translation efficiency is “TE = (the TPM_CDS_ of Ribo-seq)/(the TPM_CDS_ of RNA-seq)”. To avoid inflation due to a small denominator, only genes with an RNA-seq TPM greater than 0.5 were used in the statistical analysis of translation efficiency. The plotting of 3-nt periodicity of the Ribo-seq and coverage of RNA-seq was generated by incorporating the plot function in R v3.4.3 (78) with functions from GenomicRanges v1.30.3, GenomicFeatures v1.30.3, and GenomicAlignments v1.14.2 libraries (79) to read in the gtf file and RNA-seq bam file. The merged RNA-seq bam file from STAR and the processed "P_sites_all" file from RiboTaper were used to plot the RNA-seq coverage and P-sites of Ribo-seq, respectively. The Linux command line code to preprocess the “P_sites_all” file before used for plotting was “cut -f 1,3,6 P_sites_all | sort | uniq -c | sed -r 's/^ (*[^]+) +/\1\t/' > name_output_file”. For plotting the CUG/non-AUG start gene, the CDS range of the gene in the gtf file was manually modified before plotting.

### Statistical analysis

The statistical analysis in the paper was performed in R (78). The chisq.test and ks.test functions of the "stats" package in R were used for the Chi-squared analysis and the Kolmogorov-Smirnov test, respectively. The Pearson and Spearman correlation coefficients were calculated using the "cor" function. Pairwise comparisons were performed using the "corrplot" function in the corrplot v0.84 package (80). The empirical cumulative probabilities of translation efficiency were calculated using the "ecdf" function (in the “stats” package) and plotted with the base R plot function.

### Protein extraction and digestion

Roots (~3 cm near the tip) and shoots (shoot tip including ~1 cm hypocotyl) of four-day old tomato seedlings were harvested at ZT3 (3 hours after light on). The proteomics experiments were carried out based established methods as follows (81–83). Five volumes (v:w) of Tris buffered phenol pH 8 was added to 150 mg of ground tissue, vortexed 1 min, then mixed with 5 volumes (buffer:tissue, v:w) of extraction buffer (50 mM Tris pH 7.5, 1 mM EDTA pH 8, 0.9 M sucrose), and then centrifuge at 13,000 x g, for 10 min at 4°C. The phenol phase was transferred to a new tube and a second phenol extraction was performed on the aqueous phase. The two phenol phase extractions were combined and 5 volume of prechilled methanol with 0.1 M ammonium acetate was added. This was mixed well and keep at −80°C for 1h prior to centrifugation at 4,500 x g, for 10 min at 4°C. Precipitation with 0.1 M ammonium acetate in methanol was performed twice with incubation at −20°C for 30 min. The sample was resuspended in 70% methanol at kept at −20°C for 30 min prior to centrifuging at 4,500 x g, for 10 min at 4°C. The supernatant was discarded and the pellet was placed in a vacuum concentrator till near dry. Two volumes (buffer:pellet, v:v) of protein digestion buffer (8M urea, 50 mM Tris pH 7, 5 mM Tris(2-carboxyethyl)phosphine hydrochloride (TCEP)) was added to the pellet. The samples were then probe sonicated to aid in resuspension of the pellet. The protein concentration was then determined using the Bradford assay (Thermo Scientific).

The solubilized protein (~ 1 mg) was added to an Amicon Ultracel – 30K centrifugal filter (Cat # UFC803008) and centrifuged at 4,000 x g for 20-40 min. This step was repeated once. Then 4 ml of urea solution with 2mM TCEP was added to the filter unit and centrifuged at 4,000 x g for 20-40 min. Next, 2 ml iodoacetamide (IAM) solution (50 mM IAM in 8 M urea) was added and incubated without mixing at room temperature for 30 min in the dark prior to centrifuging at 4,000 x g for 20-40 min. Two ml of urea solution was added to the filter unit, which was then centrifuged at 4,000 x g for 20-40 min. This step was repeated once. Two ml of 0.05 M NH_4_HCO_3_ was added to the filter unit and centrifuged at 4,000 x g for 20-40 min. This step was repeated once. Then 2 ml 0.05M NH_4_HCO_3_ with trypsin (enzyme to protein ratio 1:100) or GluC (enzyme to protein ratio 1:20) was added. Samples were incubated at 37°C overnight. Undigested protein was estimated using Bradford assays then trypsin (1 μg/μl) was added to a ratio of 1:100 and a equal volume of Lys-C (0.1 μg/μl) were added to the trypsin/Lys-C digested sample and GluC was added at a ratio of 1: 20 to the sample digested with GluC. The digests were incubated for an additional 4 hours at 37°C. The filter unit was added to a new collection tube and centrifuged at 4,000 x g for 20-40 min. 1 ml 0.05M NH_4_HCO_3_ was added and centrifuged at 4,000 x g for 20-40 min. The samples were acidified to pH 2-3 with 100% formic acid and centrifuged at 21,000 x g for 20 min. Finally, samples were desalted using 50 mg Sep-Pak C18 cartridges (Waters). Eluted peptides were dried using a vacuum centrifuge (Thermo) and resuspended in 0.1% formic acid. Peptide amount was quantified using the Pierce BCA Protein assay kit.

### LC/MS-MS

An Agilent 1260 quaternary HPLC was used to deliver a flow rate of ~600 nL min-1 via a splitter. All columns were packed in house using a Next Advance pressure cell and the nanospray tips were fabricated using fused silica capillary that was pulled to a sharp tip using a laser puller (Sutter P-2000). 25 μg of peptides were loaded unto 20 cm capillary columns packed with 5 μM Zorbax SB-C18 (Agilent), which was connected using a zero dead volume 1 μm filter (Upchurch, M548) to a 5 cm long strong cation exchange (SCX) column packed with 5 μm PolySulfoethyl (PolyLC). The SCX column was then connected to a 20 cm nanospray tip packed with 2.5 μM C18 (Waters). The 3 sections were joined and mounted on a custom electrospray source for on-line nested peptide elution. A new set of columns was used for every sample. Peptides were eluted from the loading column unto the SCX column using a 0 to 80% acetonitrile (ACN) gradient over 60 minutes. Peptides were then fractionated from the SCX column using a series of ammonium acetate salt steps as following:10, 30, 32.5, 35, 37.5, 40, 42.5, 45, 50, 55, 65, 75, 85, 90, 95, 100, 150, and 1000 mM. For these analyses, buffers A (99.9% H_2_O, 0.1% formic acid), B (99.9% ACN, 0.1% formic acid), C (100 mM ammonium acetate, 2% formic acid), and D (1 M ammonium acetate, 2% formic acid) were utilized. For each salt step, a 150-minute gradient program comprised of a 0–5 minute increase to the specified ammonium acetate concentration (using buffers C or D), 5–10 minutes hold, 10–14 minutes at 100% buffer A, 15–120 minutes 5–35% buffer B, 120–140 minutes 35–80% buffer B, 140–145 minutes 80% buffer B, and 145–150 minutes buffer A was employed.

Eluted peptides were analyzed using a Thermo Scientific Q-Exactive Plus high-resolution quadrupole Orbitrap mass spectrometer, which was directly coupled to the HPLC. Data dependent acquisition was obtained using Xcalibur 4.0 software in positive ion mode with a spray voltage of 2.00 kV and a capillary temperature of 275 °C and an RF of 60. MS1 spectra were measured at a resolution of 70,000, an automatic gain control (AGC) of 3e6 with a maximum ion time of 100 ms and a mass range of 400-2000 m/z. Up to 15 MS2 were triggered at a resolution of 17,500. An AGC of 1e5 with a maximum ion time of 50 ms, an isolation window of 1.5 m/z, and a normalized collision energy of 28. Charge exclusion was set to unassigned, 1, 5–8, and >8. MS1 that triggered MS2 scans were dynamically excluded for 25s.

### Database search and FDR filtering

The raw data were analyzed using MaxQuant version 1.6.3.3 (84). A customized protein database containing 22513 proteins (Dataset_S5) was generated from the RiboTaper output file “ORFs_max_filt.” Spectra were searched against the customized protein database which was complemented with reverse decoy sequences and common contaminants by MaxQuant. Carbamidomethyl cysteine was set as a fixed modification while methionine oxidation and protein N-terminal acetylation were set as variable modifications. Digestion parameters were set to “specific” and “Trypsin/P;LysC” or “GluC”. Up to two missed cleavages were allowed. A false discovery rate less than 0.01 and protein identification level was required. The “second peptide” option was used to identify co-fragmented peptides. The “match between runs” feature of MaxQuant was not utilized. Raw data files and MaxQuant Search results have been deposited in the Mass Spectrometry Interactive Virtual Environment (MassIVE) repository: https://massive.ucsd.edu/ProteoSAFe/static/massive.jsp with dataset identifier: MSV000083363.

### Prediction of the subcellular localization of sORFs

A fasta file containing the sORF amino acid sequences was uploaded to the TargetP website (85). We selected "Plant" as the organism group and ">0.90" as the specificity cutoff and then submitted for analysis.

### Evolutionary analysis

The "tblastn" function for BLAST v2.7.1 (OS Linux_x86_64)(86) was used for the homology search. The sORFs that encoded 16~100 a.a. residues were selected for this analysis, and the reference genomes (Athaliana_167_TAIR9.fa, Atrichopoda_291_v1.0.fa, Csinensis_154_v1.fa, Mtruncatula_285_Mt4.0.fa, Osativa_323_v7.0.fa, Ppatens_318_v3.fa, S_lycopersicum_chromosomes.3.00.fa, Sitalica_312_v2.fa, Smoellendorffii_91_v1.fa, Stuberosum_448_v4.03.fa) were downloaded from Phytozome v12 (87).The fa (fasta) files for each genome were used to generate blast databases with the following code: “makeblastdb -in genome.fa -parse_seqids -dbtype nucl”, where genome.fa was replaced with the fasta file for each genome. Next, the code "tblastn -query input.fa -db species_database -out species_blast_result.txt -evalue 0.001 -outfmt '6 qseqid sseqid length qlen qstart qend sstart send pident gapopen mismatch evalue bitscore' -num_threads 10" was used to search for sequence homologs in the target genomes. The names of “species_database” and “species_blast_result.txt” were changed correspondingly. The final heatmap for a.a. identity was plotted in R using the pheatmap v1.0.10 (88) and RColorBrewer v1.1.2 libraries (89).

### miRNA target identification

The tomato miRNA sequences were extracted from Kaur *et al.* and Liu *et al.* (90, 91). Next, we used psRNATarget (51) against ITAG3.2 mRNA sequences to identify potential miRNA targets. We used “Schema V2 (2017 release)” (51) and selected “calculate target accessibility” as the analysis parameters.

### GO term analysis

agriGO v2.0 (92) was used for the GO analysis of uORF-containing genes.

## Acknowledgments

We thank Philip N. Benfey at Duke University for generous support to help initiate this project. This work used the Vincent J. Coates Genomics Sequencing Laboratory at UC Berkeley, supported by an NIH S10 OD018174 Instrumentation Grant. This research was funded by a USDA NIFA postdoctoral fellowship (award number 2016-67012-24720) and Michigan State University startup grant to PYH, and NSF (1759023), USDA NIFA Hatch project 3808, and the ISU Plant Sciences Institute awards to JWW.

## Author Contributions

HLW and PYH designed the research; PYH performed the sequencing experiments; HLW and PYH analyzed the sequencing data; GS and JWW performed the proteomic experiments and analyzed the proteomic data; HLW and PYH wrote the paper with input from all authors.

## Supplemental Information

### Dataset_S1.xlsx, spreadsheets (A) to (I)

(A): ORF_ccds (annotated ORFs) from RiboTaper output “ORF_max_filt”

(B): uORFs from RiboTaper output “ORF_max_filt”

(C): sORFs from RiboTaper output “ORF_max_filt” (D): TargetP results for sORFs

(E): sORF MassSpec spectra (F): uORF MassSpec spectra

(G): 64 ORFs using an upstream start rather than annotated start identified by RiboTaper (H): miRNAs used for psRNATarget prediction

(I): predicted miRNA-targets by psRNATarget

### Dataset_S2.xlsx: Proteogenomics, spreadsheets (A) to (C)

(A) MaxQuant_proteinGroups

(B) MaxQuant_peptides

(C) MaxQuant_modificationSpecificPeptides

**Dataset_S3_uORF.gtf** (gtf for uORFs)

**Dataset_S4_sORF.gtf** (gtf for sORFs)

**Dataset_S5.fasta** (amino acid sequences for all translated ORFs identified by RiboTaper in this study)

---

**Figure S1:**
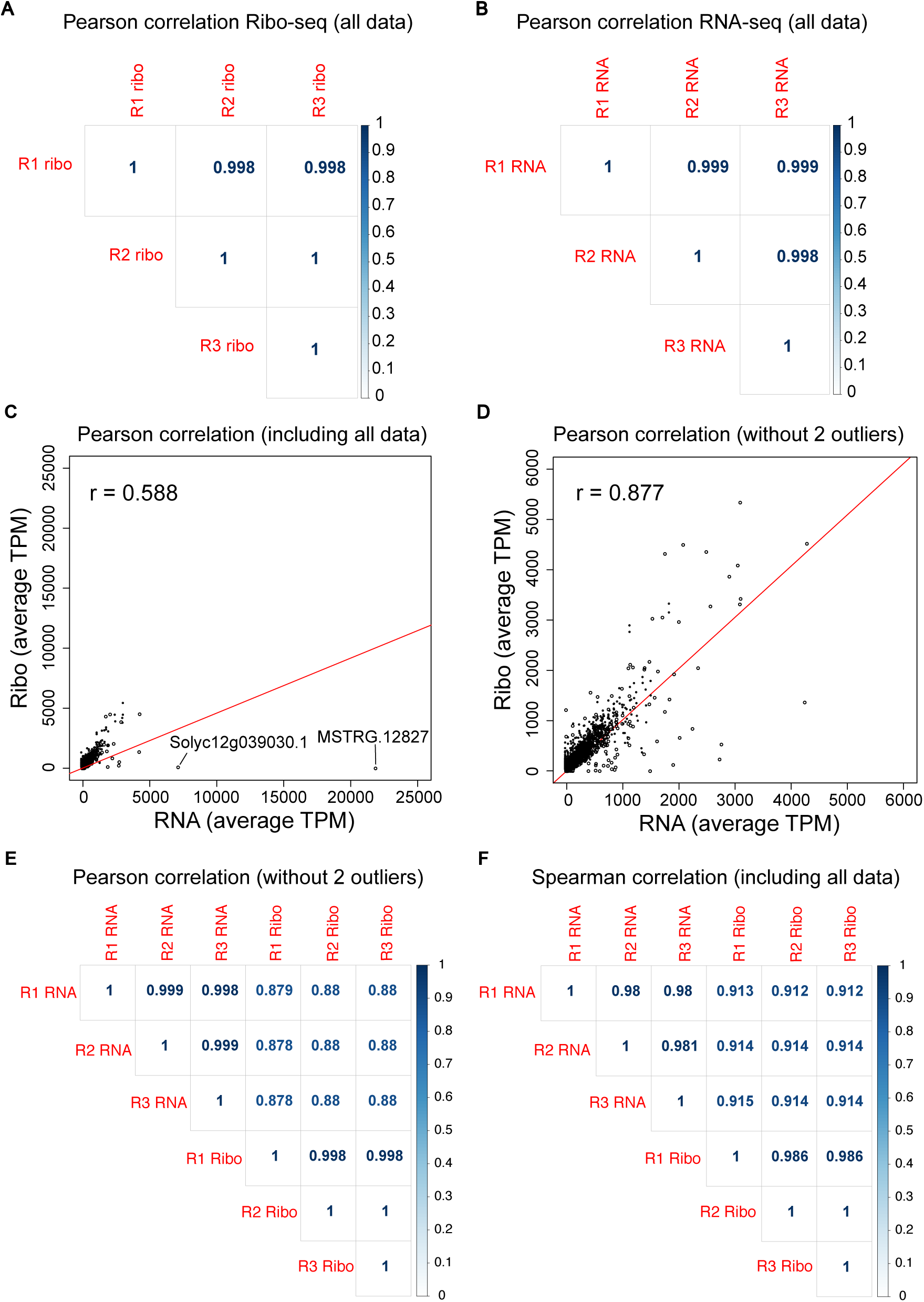
Correlation between RNA-seq and Ribo-seq data. (A-B) Pearson correlation among Ribo-seq (A) and RNA-seq (B) data from three biological replicates. (C) Pearson correlation between Ribo-seq and RNA-seq including all data points. The Pearson correlation can be strongly affected by outliers. There are two extreme outliers in our data, which strongly skew the regression line (red) and affect the correlation coefficient. (D) Same as (C) after removing the two outliers. (E) Pairwise Pearson correlation among the RNA-seq and Ribo-seq data from the three replicates after removing the two outliers. (F) Pairwise Spearman correlation among the three replicates of RNA-seq and Ribo-seq data with all data points. The Spearman method is insensitive to outliers.

**Figure S2:**
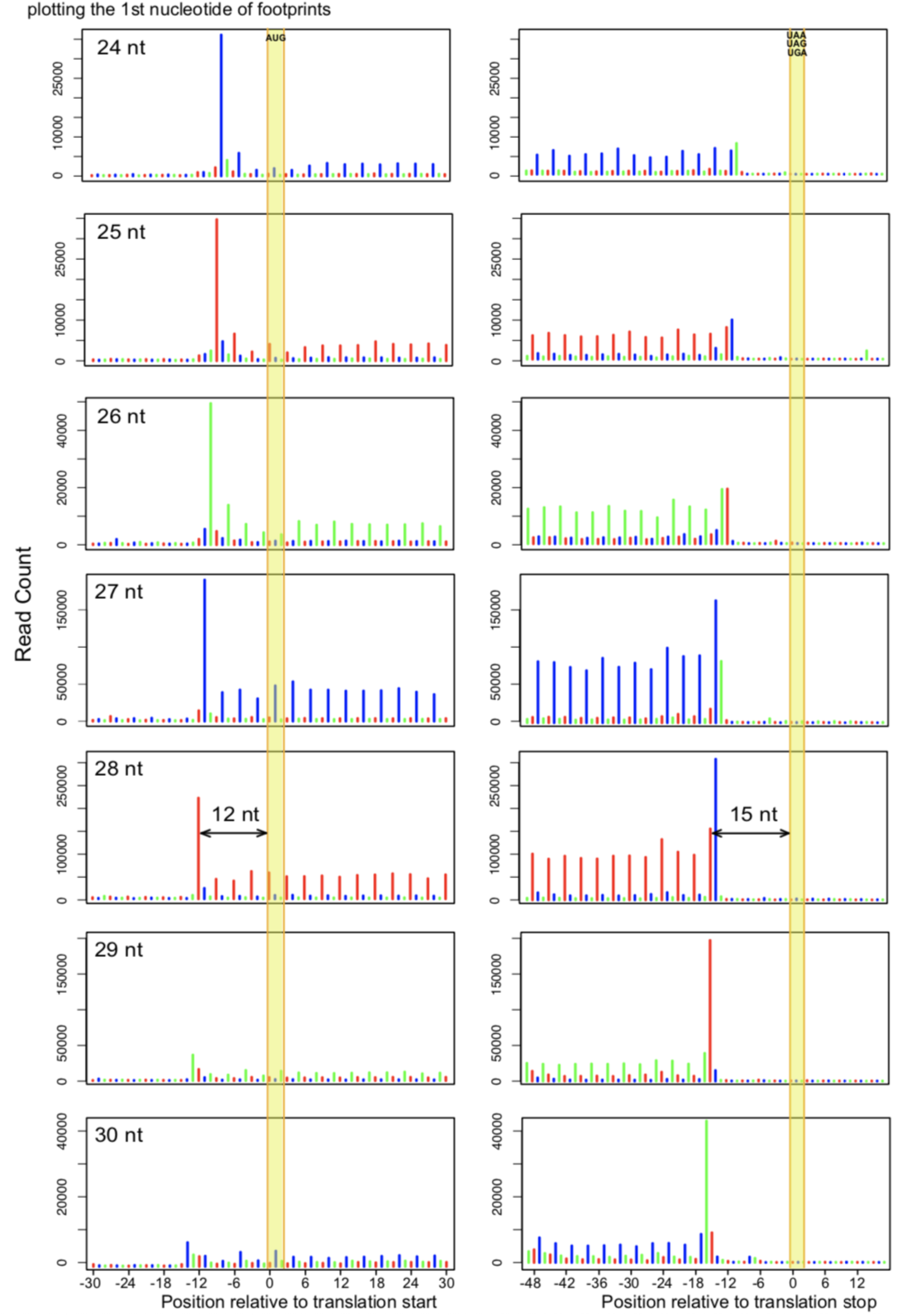
Meta-gene analysis and inference of the P-site for ribosome footprints of different lengths. Footprints near the annotated translation start (left) and translation stop (right) are shown. The position of each footprint is indicated by plotting the first nucleotide of the footprint. These metaplots were used to infer the position of the P-site in footprints of different length. For example, in 28-nt footprints, a major peak was observed the 12^th^ nt upstream of the start codon; thus, the P-site was inferred to be the 13-15^th^ nts within the footprint. Consistent with this observation, the last in-frame major peak was located at the 15^th^ nt upstream of the stop codon; thus, the A-site was inferred as the 16-18^th^ nts within the footprint.

**Figure S3:**
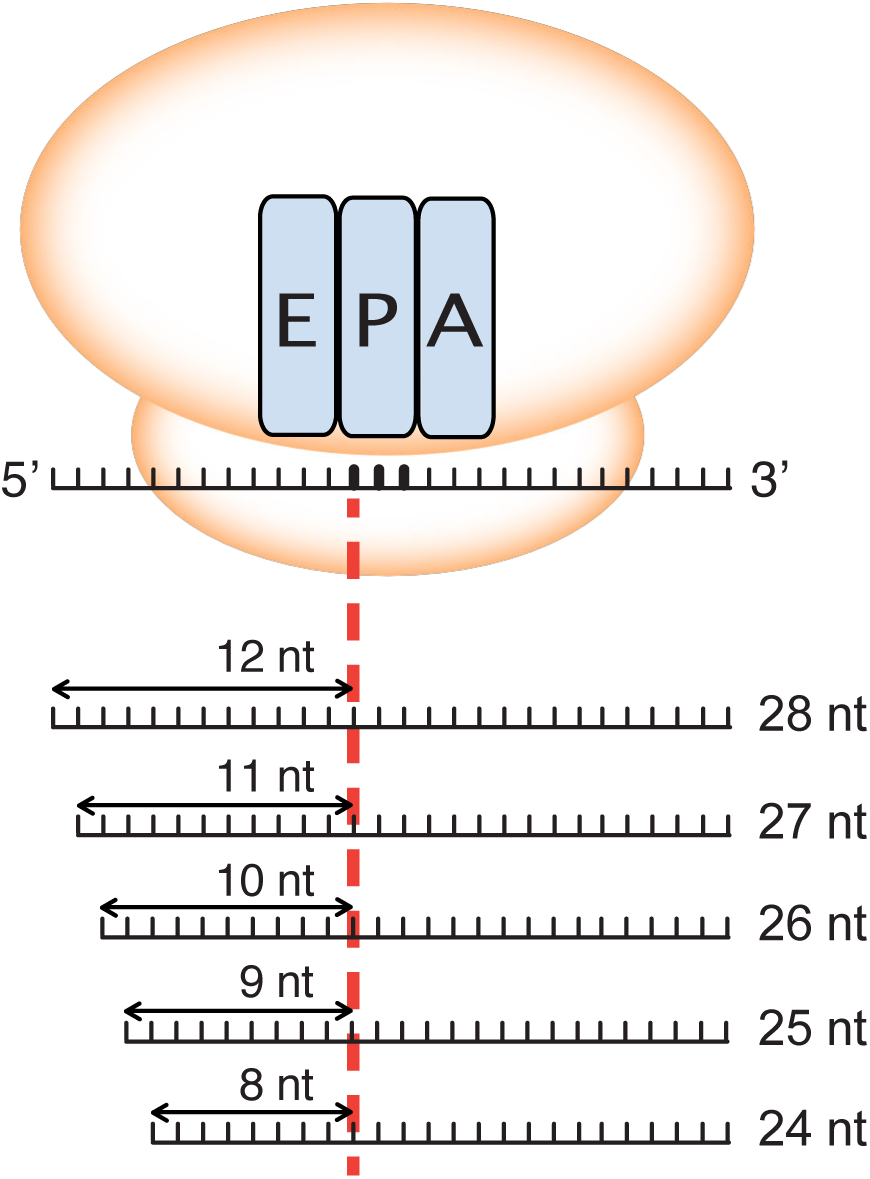
Summary of the inferred P-site position for each footprint length. The inferred P-site positions for 28-, 27-, 26-, 25- and 24-nt footprints are the 13^th^, 12^th^, 11^th^, 10^th^, and 9^th^ nts within the footprint, respectively.

**Figure S4:**
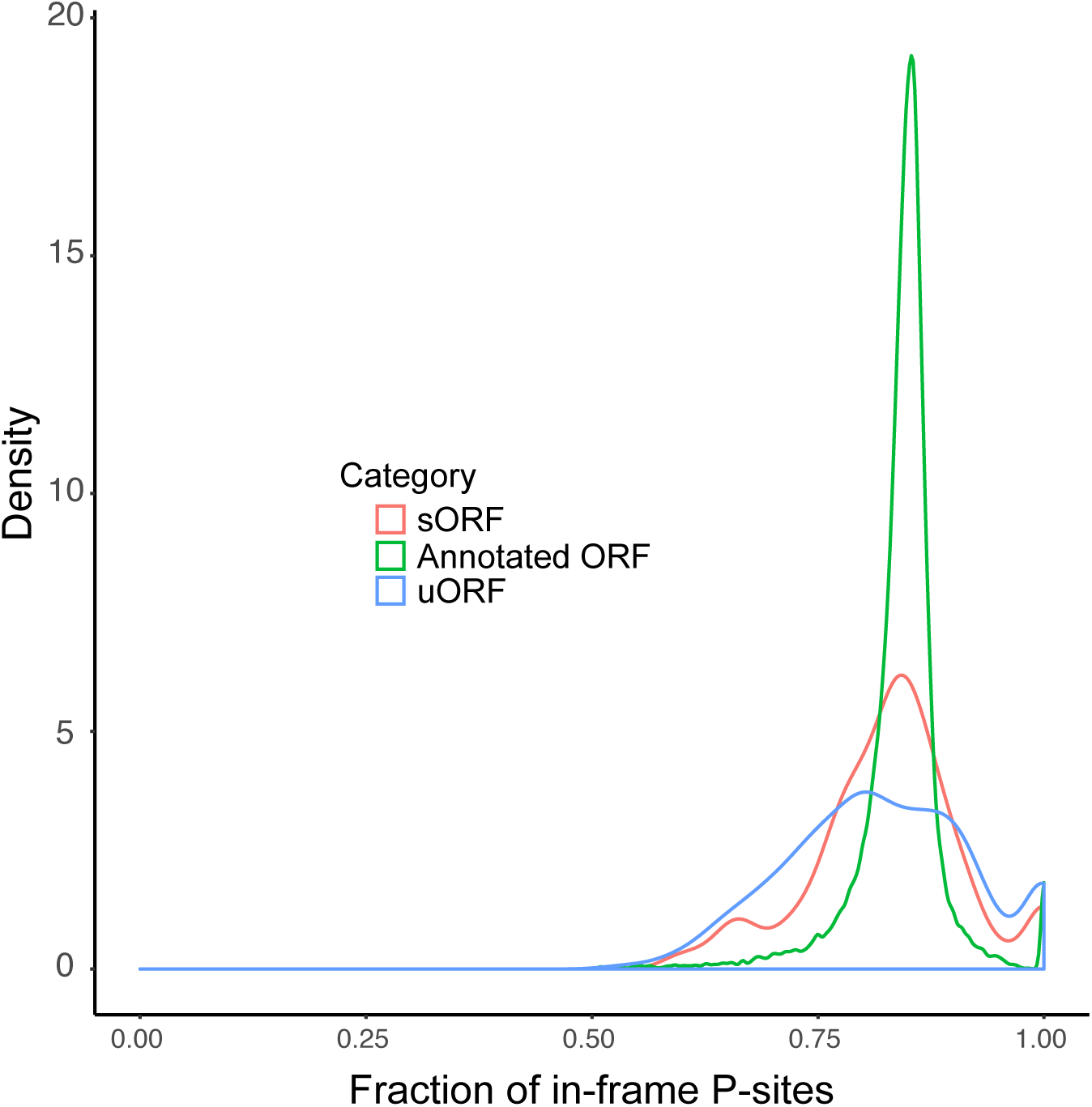
Fractions of in-frame P-sites for different groups of translated ORFs.

**Figure S5:**
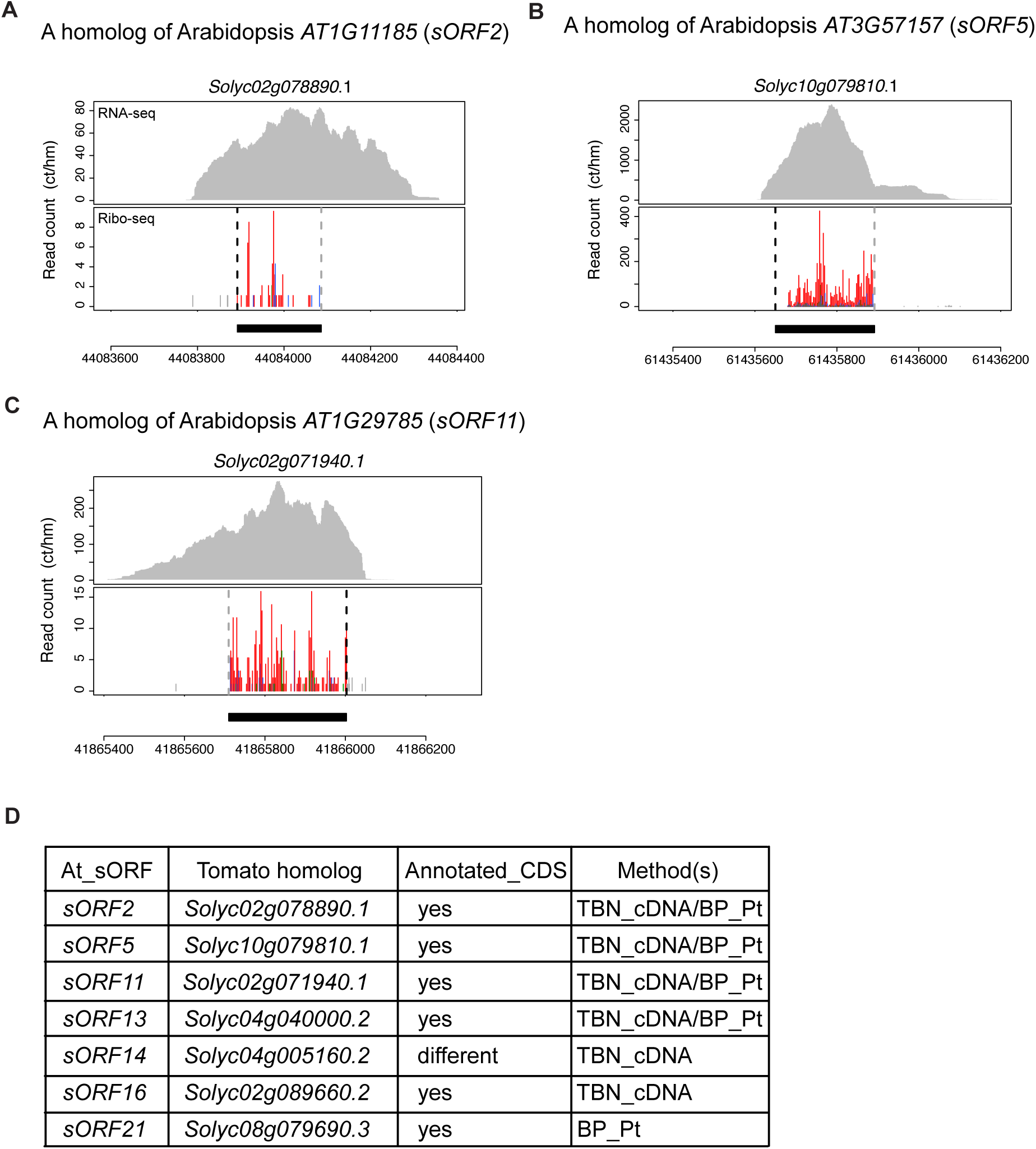
Translation of tomato homologs of Arabidopsis sORFs. (A to C) RNA-seq and Ribo-seq data for three tomato homologs of Arabidopsis sORFs. The gene models and data presentation are the same as those described in the legend of Fig. 3. (D) Seven of eight tomato homologs of the Arabidopsis sORFs were transcribed and translated in our tomato root data. The “annotated_CDS” column indicates whether the homologous sequences were identified in the annotated tomato coding regions. The “Method(s)” column indicates how the homologous sequences were identified. TBN_cDNA: tBLASTn against cDNA sequences; BP_Pt: BLASTp against protein sequences.

**Figure S6:**
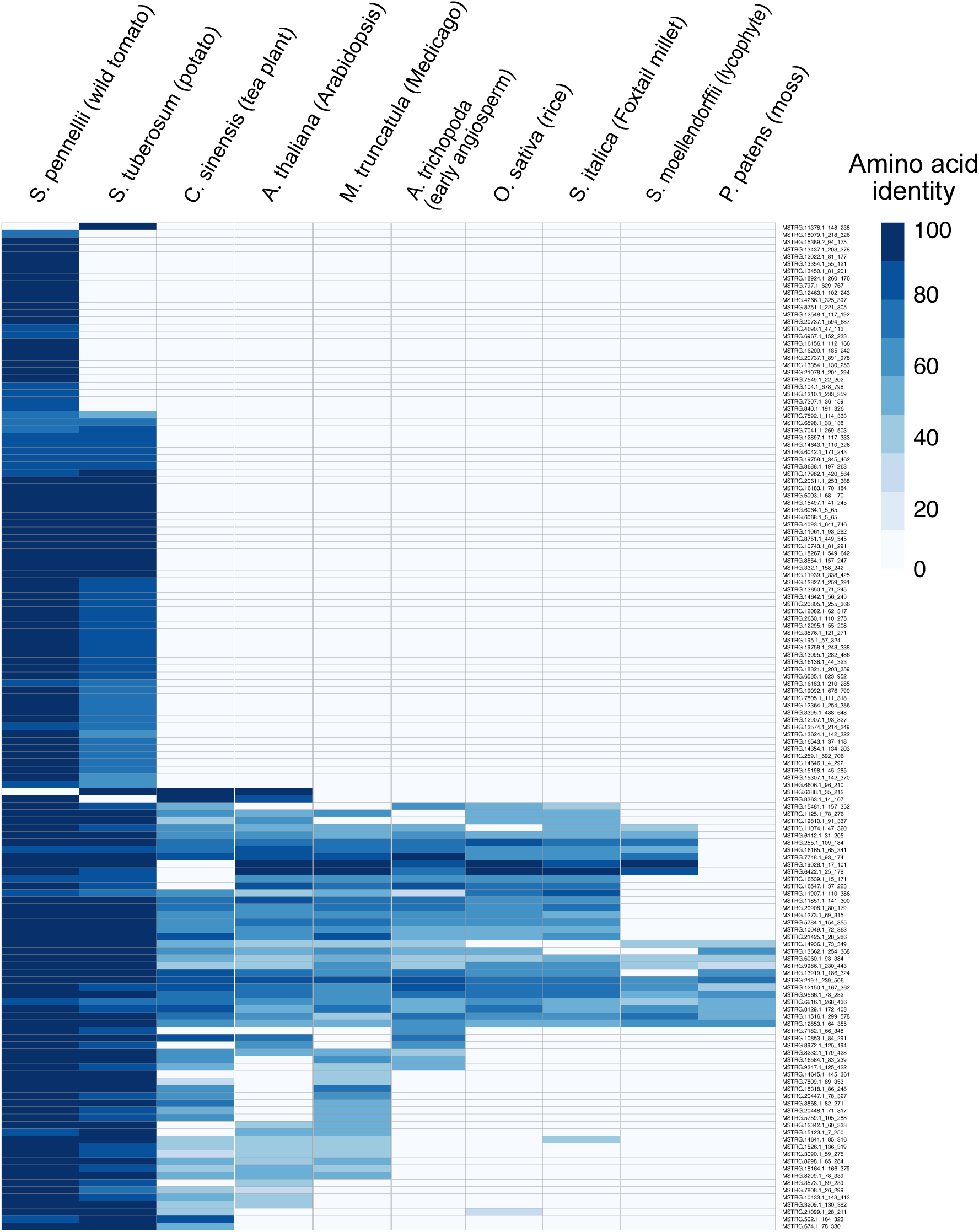
Evolutionary conservation of sORFs. Ten species, including two other species in the Solanaceae, four dicots, two monocots, a lycophyte, and a moss, were used in the analysis. The percent amino acid identity was used as an indication of sequence homology.

**Figure S7:**
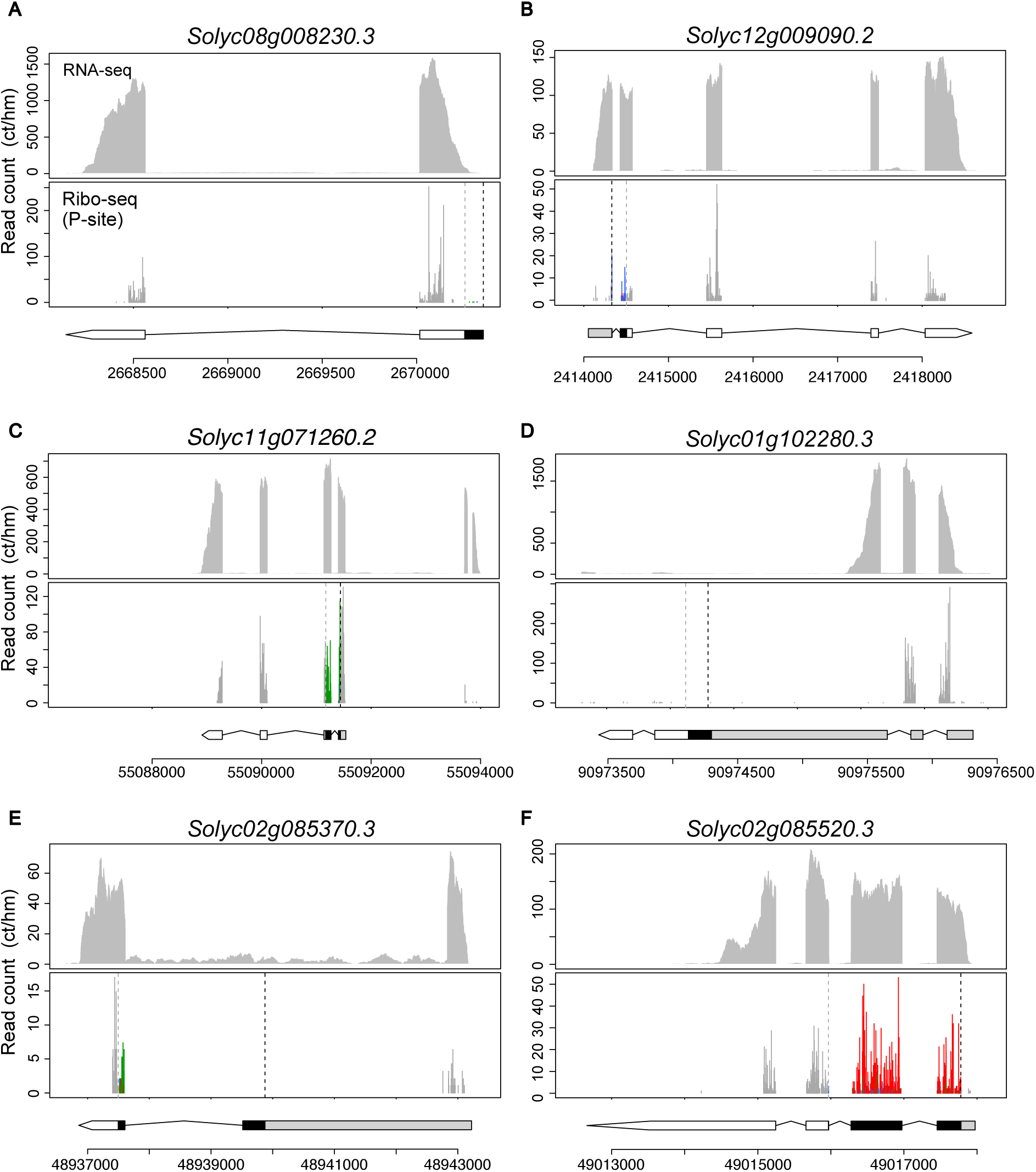
Examples of conflicts between annotated gene models and translational profiles. (A, B) Annotated ORFs do not overlap or are not in frame with the Ribo-seq reads. (C to F) Annotated isoforms are different from the expressed isoforms, and the annotated ORFs are obviously different from translated ORFs. The gene models and sequencing data presentation are the same as those described in the legend of Fig. 3.

## References

1. Sagor GHM, et al. (2016) A novel strategy to produce sweeter tomato fruits with high sugar contents by fruit-specific expression of a single bZIP transcription factor gene. Plant Biotechnol J 14(4):1116–1126.

2. Xu G, et al. (2017) uORF-mediated translation allows engineered plant disease resistance without fitness costs. Nature 545(7655):491–494.

3. Zhang H, et al. (2018) Genome editing of upstream open reading frames enables translational control in plants. Nat Biotechnol 36(9):894–898.

4. Ingolia NT, Ghaemmaghami S, Newman JRS, Weissman JS (2009) Genome-wide analysis in vivo of translation with nucleotide resolution using ribosome profiling. Science 324(5924):218–23.

5. Brar GA, Weissman JS (2015) Ribosome profiling reveals the what, when, where and how of protein synthesis. Nat Rev Mol Cell Biol 16(11):651–664.

6. Andreev DE, et al. (2017) Insights into the mechanisms of eukaryotic translation gained with ribosome profiling. Nucleic Acids Res 45(2):513–526.

7. Bazzini AA, et al. (2014) Identification of small ORFs in vertebrates using ribosome footprinting and evolutionary conservation. EMBO J 33(9):981–993.

8. Calviello L, et al. (2016) Detecting actively translated open reading frames in ribosome profiling data. Nat Methods 13:165–170.

9. Fields AP, et al. (2015) A Regression-Based Analysis of Ribosome-Profiling Data Reveals a Conserved Complexity to Mammalian Translation. Mol Cell 60(5):816–827.

10. Ji Z, Song R, Regev A, Struhl K (2015) Many lncRNAs, 5’UTRs, and pseudogenes are translated and some are likely to express functional proteins. Elife 4:e08890.

11. Hsu PY, et al. (2016) Super-resolution ribosome profiling reveals unannotated translation events in Arabidopsis. Proc Natl Acad Sci U S A 113(45):E7126–E7135.

12. Chew G-L, Pauli A, Schier AF (2016) Conservation of uORF repressiveness and sequence features in mouse, human and zebrafish. Nat Commun 7(1):11663.

13. Johnstone TG, Bazzini AA, Giraldez AJ (2016) Upstream ORFs are prevalent translational repressors in vertebrates. EMBO J:e201592759.

14. Brar GA, et al. (2012) High-resolution view of the yeast meiotic program revealed by ribosome profiling. Science 335(6068):552–7.

15. Liu M-J, et al. (2013) Translational landscape of photomorphogenic Arabidopsis. Plant Cell 25(10):3699–710.

16. Hsu PY, Benfey PN (2018) Small but Mighty: Functional Peptides Encoded by Small ORFs in Plants. Proteomics 18(10):1700038.

17. von Arnim AG, Jia Q, Vaughn JN (2013) Regulation of plant translation by upstream open reading frames. Plant Sci 214:1–12.

18. Bazin J, et al. (2017) Global analysis of ribosome-associated noncoding RNAs unveils new modes of translational regulation. Proc Natl Acad Sci U S A 114(46):E10018–E10027.

19. Ruiz-Orera J, Albà MM (2018) Translation of Small Open Reading Frames: Roles in Regulation and Evolutionary Innovation. Trends Genet. doi:10.1016/j.tig.2018.12.003.

20. Basrai MA, Hieter P, Boeke JD (1997) Small Open Reading Frames: Beautiful Needles in the Haystack. Genome Res 7(8):768–771.

21. Claverie J (1997) Computational methods for the identification of genes in vertebrate genomic sequences. Hum Mol Genet 6(10):1735–1744.

22. Zoschke R, Watkins KP, Barkan A (2013) A rapid ribosome profiling method elucidates chloroplast ribosome behavior in vivo. Plant Cell 25(6):2265–75.

23. Chotewutmontri P, Barkan A (2016) Dynamics of Chloroplast Translation during Chloroplast Differentiation in Maize. PLOS Genet 12(7):e1006106.

24. Juntawong P, Girke T, Bazin J, Bailey-Serres J (2014) Translational dynamics revealed by genome-wide profiling of ribosome footprints in Arabidopsis. Proc Natl Acad Sci U S A 111(1):E203–12.

25. Merchante C, et al. (2015) Gene-Specific Translation Regulation Mediated by the Hormone-Signaling Molecule EIN2. Cell 163(3):684–697.

26. Lei L, et al. (2015) Ribosome profiling reveals dynamic translational landscape in maize seedlings under drought stress. Plant J 84(6):1206–18.

27. Xu G, et al. (2017) Global translational reprogramming is a fundamental layer of immune regulation in plants. Nature 545(7655):487–490.

28. Shamimuzzaman M, Vodkin L (2018) Ribosome profiling reveals changes in translational status of soybean transcripts during immature cotyledon development. PLoS One 13(3):e0194596.

29. Li S, et al. (2016) Biogenesis of phased siRNAs on membrane-bound polysomes in Arabidopsis. Elife 5:e22750.

30. Schwarz D, Thompson AJ, Kläring H-P (2014) Guidelines to use tomato in experiments with a controlled environment. Front Plant Sci 5:625.

31. Fernandez-Pozo N, et al. (2015) The Sol Genomics Network (SGN)—from genotype to phenotype to breeding. Nucleic Acids Res 43(D1):D1036–D1041.

32. Guydosh NR, Green R (2014) Dom34 rescues ribosomes in 3’ untranslated regions. Cell 156(5):950–62.

33. Chung BY, et al. (2015) The use of duplex-specific nuclease in ribosome profiling and a user-friendly software package for Ribo-seq data analysis. RNA 10:1731–45.

34. Schafer S, et al. (2015) Translational regulation shapes the molecular landscape of complex disease phenotypes. Nat Commun 6:7200.

35. Pertea M, et al. (2015) StringTie enables improved reconstruction of a transcriptome from RNA-seq reads. Nat Biotechnol 33(3):290–295.

36. Pertea M, Kim D, Pertea GM, Leek JT, Salzberg SL (2016) Transcript-level expression analysis of RNA-seq experiments with HISAT, StringTie and Ballgown. Nat Protoc 11(9):1650–1667.

37. Ku HM, Vision T, Liu J, Tanksley SD (2000) Comparing sequenced segments of the tomato and Arabidopsis genomes: large-scale duplication followed by selective gene loss creates a network of synteny. Proc Natl Acad Sci U S A 97(16):9121–6.

38. Walley JW, Briggs SP (2015) Dual use of peptide mass spectra: Protein atlas and genome annotation. Curr plant Biol 2:21–24.

39. Szymanski J, et al. (2017) Label-free deep shotgun proteomics reveals protein dynamics during tomato fruit tissues development. Plant J 90(2):396–417.

40. Mata CI, et al. (2017) In-depth characterization of the tomato fruit pericarp proteome. Proteomics 17(1–2):1600406.

41. Michel AM, et al. (2012) Observation of dually decoded regions of the human genome using ribosome profiling data. Genome Res 22(11):2219–29.

42. Kearse MG, Wilusz JE (2017) Non-AUG translation: a new start for protein synthesis in eukaryotes. Genes Dev 31(17):1717–1731.

43. Simpson GG, et al. (2010) Noncanonical translation initiation of the Arabidopsis flowering time and alternative polyadenylation regulator FCA. Plant Cell 22(11):3764–77.

44. Spealman P, et al. (2018) Conserved non-AUG uORFs revealed by a novel regression analysis of ribosome profiling data. Genome Res 28(2):214–222.

45. Laing WA, et al. (2015) An upstream open reading frame is essential for feedback regulation of ascorbate biosynthesis in Arabidopsis. Plant Cell 27(3):772–86.

46. Imai A, et al. (2006) The dwarf phenotype of the Arabidopsis acl5 mutant is suppressed by a mutation in an upstream ORF of a bHLH gene. Development 133(18):3575–85.

47. Kozak M (1987) An analysis of 5’-noncoding sequences from 699 vertebrate messenger RNAs. Nucleic Acids Res 15(20):8125–48.

48. Lütcke HA, et al. (1987) Selection of AUG initiation codons differs in plants and animals. EMBO J 6(1):43–8.

49. Yu Y, Jia T, Chen X (2017) The ‘how’ and ‘where’ of plant microRNAs. New Phytol 216(4):1002–1017.

50. Li Z, Xu R, Li N (2018) MicroRNAs from plants to animals, do they define a new messenger for communication? Nutr Metab (Lond) 15(1):68.

51. Dai X, Zhuang Z, Zhao PX (2018) psRNATarget: a plant small RNA target analysis server (2017 release). Nucleic Acids Res 46(W1):W49–W54.

52. Ali Faghihi M, Wahlestedt C (2009) Regulatory roles of natural antisense transcripts doi:10.1038/nrm2738.

53. Young R, Bremer H (1976) Polypeptide-chain-elongation rate in Escherichia coli B/r as a function of growth rate. Biochem J 160(2):185–94.

54. Wu B, Eliscovich C, Yoon YJ, Singer RH (2016) Translation dynamics of single mRNAs in live cells and neurons. Science (80-) 352(6292):1430–1435.

55. Ingolia NT (2016) Ribosome Footprint Profiling of Translation throughout the Genome. Cell 165(1):22–33.

56. Lease KA, Walker JC (2006) The Arabidopsis unannotated secreted peptide database, a resource for plant peptidomics. Plant Physiol 142(3):831–8.

57. Hanada K, Zhang X, Borevitz JO, Li W-H, Shiu S-H (2007) A large number of novel coding small open reading frames in the intergenic regions of the Arabidopsis thaliana genome are transcribed and/or under purifying selection. Genome Res 17(5):632–640.

58. Hanada K, et al. (2010) sORF finder: a program package to identify small open reading frames with high coding potential. Bioinformatics 26(3):399–400.

59. Hanada K, et al. (2013) Small open reading frames associated with morphogenesis are hidden in plant genomes. Proc Natl Acad Sci U S A 110(6):2395–400.

60. Ghorbani S, et al. (2015) Expanding the repertoire of secretory peptides controlling root development with comparative genome analysis and functional assays. J Exp Bot 66(17):5257–69.

61. Tavormina P, De Coninck B, Nikonorova N, De Smet I, Cammue BPA (2015) The Plant Peptidome: An Expanding Repertoire of Structural Features and Biological Functions. Plant Cell 27(8):2095–118.

62. Berardini TZ, et al. (2015) The Arabidopsis information resource: Making and mining the “gold standard” annotated reference plant genome. Genesis 53(8):474–85.

63. Crappe J, et al. (2014) PROTEOFORMER: deep proteome coverage through ribosome profiling and MS integration. Nucleic Acids Res 43(5):1–10.

64. Menschaert G, et al. (2013) Deep proteome coverage based on ribosome profiling aids mass spectrometry-based protein and peptide discovery and provides evidence of alternative translation products and near-cognate translation initiation events. Mol Cell Proteomics 12(7):1780–90.

65. Van Damme P, Gawron D, Van Criekinge W, Menschaert G (2014) N-terminal proteomics and ribosome profiling provide a comprehensive view of the alternative translation initiation landscape in mice and men. Mol Cell Proteomics 13(5):1245–61.

66. Ruggles K V, et al. (2017) Methods, Tools and Current Perspectives in Proteogenomics. Mol Cell Proteomics 16(6):959–981.

67. Agrawal GK, Jwa N-S, Lebrun M-H, Job D, Rakwal R (2010) Plant secretome: Unlocking secrets of the secreted proteins. Proteomics 10(4):799–827.

68. Casson SA (2002) The POLARIS Gene of Arabidopsis Encodes a Predicted Peptide Required for Correct Root Growth and Leaf Vascular Patterning. PLANT CELL ONLINE 14(8):1705–1721.

69. Blanvillain R, et al. (2011) The Arabidopsis peptide kiss of death is an inducer of programmed cell death. EMBO J 30(6):1173–1183.

70. Valdivia ER, et al. (2012) DVL genes play a role in the coordination of socket cell recruitment and differentiation. J Exp Bot 63(3):1405–1412.

71. Ikeuchi M, et al. (2011) ROTUNDIFOLIA4 Regulates Cell Proliferation Along the Body Axis in Arabidopsis Shoot. Plant Cell Physiol 52(1):59–69.

72. De Coninck B, et al. (2013) Mining the genome of Arabidopsis thaliana as a basis for the identification of novel bioactive peptides involved in oxidative stress tolerance. J Exp Bot 64(17):5297–5307.

73. Langmead B, Salzberg SL (2012) Fast gapped-read alignment with Bowtie 2. Nat Methods 9(4):357–9.

74. Liao Y, Smyth GK, Shi W (2014) featureCounts: an efficient general purpose program for assigning sequence reads to genomic features. Bioinformatics 30(7):923–930.

75. Dobin A, et al. (2013) STAR: ultrafast universal RNA-seq aligner. Bioinformatics 29(1):15–21.

76. Li H, et al. (2009) The Sequence Alignment/Map format and SAMtools. Bioinformatics 25(16):2078–2079.

77. Li B, Dewey CN (2011) RSEM: accurate transcript quantification from RNA-Seq data with or without a reference genome. BMC Bioinformatics 12(1):323.

78. R Core Team (2017) (2017) R: A language and environment for statistical computing. R Found Stat Comput Vienna, Austria:R Foundation for Statistical Computing.

79. Lawrence M, et al. (2013) Software for computing and annotating genomic ranges. PLoS Comput Biol 9(8):e1003118.

80. Wei T (2013) corrplot: Visualization of a correlation matrix. Available at: https://cran.r-project.org/web/packages/corrplot/index.html [Accessed April 27, 2016].

81. Castellana NE, et al. (2014) An automated proteogenomic method uses mass spectrometry to reveal novel genes in Zea mays. Mol Cell Proteomics 13(1):157–67.

82. Song G, Hsu PY, Walley JW (2018) Assessment and Refinement of Sample Preparation Methods for Deep and Quantitative Plant Proteome Profiling. Proteomics 18(17):1800220.

83. Song G, Brachova L, Nikolau BJ, Jones AM, Walley JW (2018) Heterotrimeric G-Protein-Dependent Proteome and Phosphoproteome in Unstimulated Arabidopsis Roots. Proteomics 18(24):1800323.

84. Tyanova S, Temu T, Cox J (2016) The MaxQuant computational platform for mass spectrometry-based shotgun proteomics. Nat Protoc 11(12):2301–2319.

85. Emanuelsson O, Nielsen H, Brunak S, von Heijne G (2000) Predicting Subcellular Localization of Proteins Based on their N-terminal Amino Acid Sequence. J Mol Biol 300(4):1005–1016.

86. Camacho C, et al. (2009) BLAST+: architecture and applications. BMC Bioinformatics 10(1):421.

87. Goodstein DM, et al. (2012) Phytozome: a comparative platform for green plant genomics. Nucleic Acids Res 40:D1178–86.

88. Kolde R (2015) pheatmap: Pretty Heatmaps. Available at: https://cran.r-project.org/web/packages/pheatmap/index.html.

89. Neuwirth E (2014) RColorBrewer: ColorBrewer Palettes.

90. Kaur P, et al. (2017) Genome-wide identification and characterization of miRNAome from tomato (Solanum lycopersicum) roots and root-knot nematode (Meloidogyne incognita) during susceptible interaction. PLoS One 12(4):e0175178.

91. Liu M, et al. (2017) Profiling of drought-responsive microRNA and mRNA in tomato using high-throughput sequencing. BMC Genomics 18(1):481.

92. Tian T, et al. (2017) agriGO v2.0: a GO analysis toolkit for the agricultural community, 2017 update. Nucleic Acids Res 45(W1):W122–W129.

